# Extreme in Every Way: Exceedingly Low Genetic Diversity in Snow Leopards Due to Persistently Small Population Size

**DOI:** 10.1101/2023.12.14.571340

**Authors:** Katherine A. Solari, Simon Morgan, Andrey D. Poyarkov, Byron Weckworth, Gustaf Samelius, Koustubh Sharma, Stephane Ostrowski, Uma Ramakrishnan, Zairbek Kubanychbekov, Shannon Kachel, Örjan Johansson, Purevjav Lkhagvajav, Heather Hemmingmoore, Dmitry Y. Alexandrov, Munkhtsog Bayaraa, Alexey Grachev, Miroslav P. Korablev, Jose A. Hernandez-Blanco, Bariushaa Munkhtsog, Barry Rosenbaum, Viatcheslav V. Rozhnov, Ali Madad Rajabi, Hafizullah Noori, Kulbhushansingh Suryawanshi, Ellie E. Armstrong, Dmitri A. Petrov

**Affiliations:** Department of Biology, Stanford University, Stanford, CA, USA; A.N. Severtsov Institute of Ecology and Evolution, Russian Academy of Science, Moscow, Russia; Panthera, New York, NY, USA; Snow Leopard Trust, Seattle, WA, USA; Nordens Ark, Hunnebostrand, Sweden; Snow Leopard Foundation in Kyrgyzstan, Bishkek, Kyrgyz Republic; Wildlife Conservation Society, New York, NY, USA; National Centre for Biological Sciences, TIFR, Bangalore, India; Ilbirs Foundation, Bishkek, Kyrgyzstan; Grimsö Wildlife Research Station, Swedish University of Agricultural Sciences, Riddarhyttan, Sweden; Snow Leopard Conservation Foundation, Ulanbaatar, Mongolia; Wildlife Institute, College of Nature Conservation of the Beijing Forestry University, Beijing, China; Institute of Zoology of Republic of Kazakhstan, Almaty, Kazakhstan; Institute of General and Experimental Biology, Academy of Sciences of Mongolia, Ulaanbaatar, Mongolia; Altai Institute for Research and Conservation, Boulder, CO, USA; Wildlife Conservation Society, Afghanistan Program, Kabul, Afghanistan; Nature Conservation Foundation, Mysore, India; Canadian Institute for Advanced Research, Toronto, Ontario, Canada; Washington State University, Pullman, WA, USA; Chan Zuckerberg BioHub, San Francisco, CA, USA; Stanford Cancer Institute, Stanford University School of Medicine, Stanford, CA, USA

## Abstract

Snow leopards (*Panthera uncia*) serve as an umbrella species whose conservation benefits their high-elevation Asian habitat. Their numbers are believed to be in decline due to numerous Anthropogenic threats; however, their conservation is hindered by numerous knowledge gaps. They are the least studied genetically of all big cat species with more to learn regarding their population structure, historical population size, and current levels of genetic diversity. Here, we use whole-genome sequencing data for 41 snow leopards (37 newly sequenced) to offer new insights into these unresolved questions. Among our samples, we find evidence of a primary genetic divide between the northern and southern part of the range around the Dzungarian Basin, as previously identified, and a secondary divide south of Kyrgyzstan around the Taklamakan Desert. Most noteworthy, we find that snow leopards have the lowest genetic diversity of any big cat species, due to a persistently small population size (relative to other big cat species) throughout their evolutionary history rather than recent inbreeding. Without a large population size or ample standing genetic variation to help buffer them from any forthcoming Anthropogenic challenges, snow leopard persistence may be more tenuous than currently appreciated.

## Introduction

Residing in some of the most extreme and remote areas of the world, snow leopards (*Panthera uncia*) are rarely encountered and are challenging to study, making them one of the most enigmatic of the large charismatic mammals. They are among the largest carnivores in the high-elevation habitat in which they reside and their persistence relies on healthy mountain ungulate populations (Jumabay-Uulu et al. 2014) sometimes supplemented by livestock (Shehzad et al. 2012; Wegge et al. 2012; Aryal et al. 2014; Johansson et al. 2015; Chetri et al. 2017; Suryawanshi et al. 2017; Lu et al. 2021; Thapa et al. 2021). Snow leopard habitat consists of mountainous areas of Asia, spanning 12 countries (Fig. 1A), habitat that offers immense ecosystem services–acting as an important source of carbon storage (Mu et al. 2020) and providing water to almost 2 billion people. Snow leopards serve as an umbrella species whose conservation benefits this globally crucial Asian mountain ecosystem. In spite of the global benefits of snow leopard conservation, efforts are often impeded by the many knowledge gaps regarding this elusive species (McCarthy and Mallon 2016).

**Figure 1.**
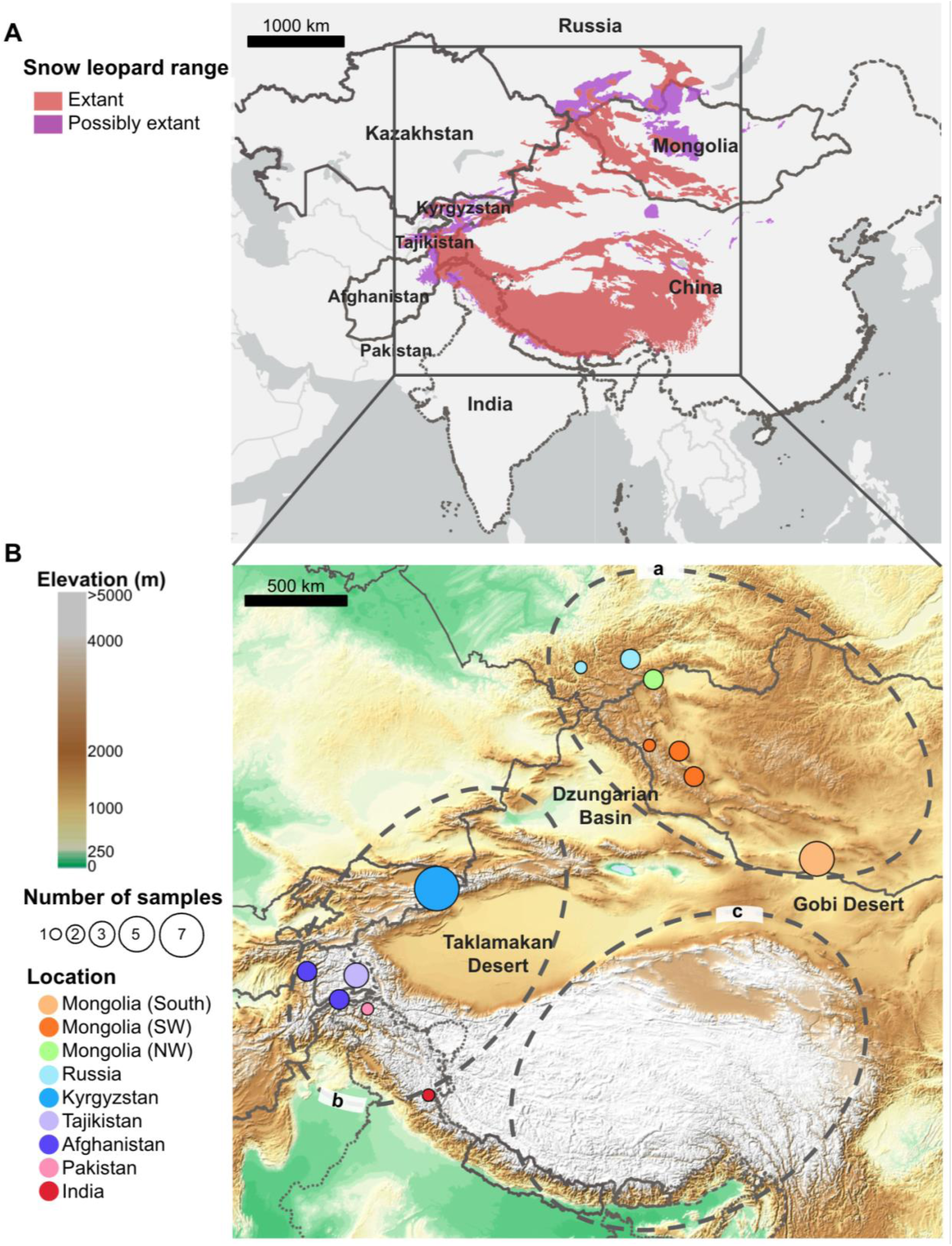
Snow leopard distribution and sample maps. A) The IUCN snow leopard distribution (McCarthy et al. 2017) is shown and the largest snow leopard range countries are labeled. B) Sample locations are indicated with different sized circles indicating the number of samples from each location. The basemap (ESRI 2013) indicates elevation and landscape features discussed in the text are labeled. Grey dashed ovals indicate the geographic distribution of the three subspecies suggested by Janecka et al. 2017–a) *P. u. irbis*, b) *P. u. uncia*, c) *P. u. uncioides*. In both maps, country boundaries available from the Snow Leopard Trust (The Snow Leopard Trust) are shown in dark grey. Not all country boundaries are in agreement, so a dotted line is used for India and a dashed line is used for China to make overlapping boundaries visible. Maps were created using ArcGIS software by ESRI.

Snow leopards were listed as Endangered by the International Union for Conservation of Nature (IUCN) for 45 years but were downlisted to Vulnerable in 2017 due to not meeting specific criteria of population size (fewer than 2,500 mature individuals) and percent population decline (more than 20% over two generations) for Endangered status. This change of status has been controversial (Aryal 2017; Ale and Mishra 2018) as snow leopard numbers are presumed to be in decline due to habitat loss, decreased availability of primary prey (high-elevation, mountain-dwelling ungulates), retaliatory killings for livestock predation (Jackson et al. 2010), and poaching for their skins (Network 2014). As climate change in high mountain Asia is occurring at an even more rapid rate than elsewhere in the Northern hemisphere, excluding the Arctic (Lalande et al. 2021), it is also likely to become an increasing threat to snow leopards (Li et al. 2016).

Currently, the global snow leopard population size is estimated to be anywhere from 4,700 to 7,500 individuals and little is known about their historical population size and range (McCarthy et al. 2024) or their current population trends (McCarthy et al. 2017). While many other big cat species experienced historical declines due to range contractions during the Last Glacial Maximum (Cooper et al. 2016; Cooper et al. 2021) and are facing a second, contemporary human-driven decline (Dobrynin et al. 2015; Pečnerová et al. 2021), it remains unclear if the snow leopard was ever more abundant than is currently estimated. One study using fecal microsatellite data suggests that snow leopards may have undergone a bottleneck ∼8,000 years ago (Janečka et al. 2017). Estimates of genetic diversity and levels of inbreeding are also limited due to a dearth of genomic data for the species. They are the least studied genetically of all the big cat species (Weckworth 2021) with whole-genome sequencing (WGS) data available for only two wild snow leopards (Cho et al. 2013) and two captive individuals (Armstrong et al. 2022) prior to this study. Genetic diversity assessed using fecal microsatellite data (Janečka et al. 2017; Aruge et al. 2019; Atzeni et al. 2021; Korablev et al. 2021; Hacker et al. 2023) as well as genomic data from one of the previously sequenced wild snow leopards suggests a lack of diversity (Cho et al. 2013), but genomic data from additional samples are required to determine if this is a characteristic of the species across its range.

Additionally, there is still more to learn about snow leopard population structure and connectivity across their range (McCarthy et al. 2016). Numerous studies have used microsatellite markers from fecal samples to assess genetic structure and connectivity at varying geographic scales. Such studies have found connectivity at the local scale to be location dependent, with evidence of continuous habitat connectivity across at least 75 km in Pakistan (Aruge et al. 2019) and >1000 km in Mongolia (Hacker et al. 2023), weak genetic differentiation among snow leopards samples across 400 km of mountainous terrain in Northern China (Atzeni et al. 2021), and signs of genetic structure between sampling areas in Russia about 500 km apart (Korablev et al. 2021). At a larger geographic scale, fecal microsatellite studies (Janečka et al. 2017; Korablev et al. 2021; Hacker et al. 2023) and other lines of evidence (Riordan et al. 2016; Li et al. 2020) suggest a geographic divide between Mongolia/Russia and the Southern part of the range due to the Dzungarian Basin and Gobi Desert. However, snow leopards are known to cross long distances between mountain ranges (McCarthy et al. 2005; Sharma et al. 2014). In addition to this prominent north-south divide, the largest scale fecal microsatellites study also identified a second divide within the southern group separating the east and west of the Himalayas-Tibetan Plateau complex (Janečka et al. 2017). Janecka et al. (Janečka et al. 2017) argue that each of these three distinct groups constitute unique subspecies; however, this subspecies demarcation and the level of connectivity across the landscape remains controversial (Janečka et al. 2018; Senn et al. 2018). Population structure and connectivity have never been assessed in snow leopards using whole-genome data.

In addition to the wild snow leopard population, the international community has worked for decades to establish a sustainable zoo population. As of 2008 there were 445 snow leopards across 205 institutions globally, not including China, representing the genetic diversity of 56 wild founders (Blomqvist 2008), most of which came from the wild in the 1960s-1990s, often from unknown locations. As it is the goal of zoos to maintain a genetically diverse population (Tetzloff and Schwartz 2024) and to act as reserves of genetic diversity for endangered species, it is important to know what portion of the global genomic diversity of snow leopards this population represents.

Here, we generate WGS data for 33 wild snow leopards from multiple locations across their range in addition to four captive snow leopards. We combine this novel data with existing data for four individuals to achieve three main objectives: 1) characterize snow leopard population structure and connectivity to see how estimates from WGS data compare to fecal microsatellite data; 2) assess the current level of genomic diversity in snow leopards, how this compares to other big cat species, and how this relates to historical population size and inbreeding levels; and 3) assess the ancestry of the current zoo snow leopard population. Among our samples, which do not include most of the central/southern part of the range, we find evidence of a genetic divide between the north and south, as well as a divide between the Kyrgyz population and populations farther south. However, low levels of genetic differentiation between groups suggests recent separation and/or weak levels of connectivity. We also find the current zoo population to be dominated by Kyrgyzstan-region ancestry. Most notably, we find snow leopards to be the least genetically diverse contemporary big cat species, owed to a persistently small population throughout their evolutionary history rather than to recent inbreeding. We believe these results have significant implications for snow leopard conservation.

## Results

We generated WGS data for 37 snow leopards–33 wild born snow leopards representing six countries (Mongolia, Russia, Kyrgyzstan, Tajikistan, Afghanistan, and Pakistan) and four from the American zoo population. After removing samples based on sequence quality and breadth we were left with 34 samples from this newly generated sample set with an average depth of coverage of 7.3X (minimum of 3X and maximum of 16.8X). We also included all published WGS data for snow leopards in our analysis, which consisted of data for two additional wild samples, one from Mongolia (Cho et al. 2013) and one from India, and one captive individual from the American zoo population (Armstrong et al. 2022) with an average depth of coverage of 23X (minimum of 12X and maximum of 28.9X). Published sequencing data for an additional captive individual was not included due to sequencing quality issues. After removing samples based on sequence quality and breadth, we were left with 37 samples–34 from our newly generated data and three previously published samples. This final dataset consisted of 32 wild born snow leopards representing all seven countries included in the pre-filtered sample set (Mongolia, Russia, Kyrgyzstan, Tajikistan, Afghanistan, Pakistan, and India) (Fig. 1B) and five captive samples (Supplementary Table 1).

We mapped these data to the snow leopard reference genome (NCBI accession PRJNA602938) (Armstrong et al. 2022) and called single nucleotide polymorphisms (SNPs) as described in the methods, resulting in a final SNP set of 1,591,978. Within this final dataset, we identified one pair of first-degree relatives and three pairs of second-degree relatives (all within the wild samples). We removed one representative from each of these four related pairs before conducting analyses that could be impacted by the presence of related individuals as indicated in the methods.

### Population structure and dispersal barriers

We assessed population structure among our samples using principal component analysis (PCA). PCA results indicated that the one Indian sample, and to a lesser extent, one of the Tajikistan samples (U13) were genetically distinct from all of the other samples (Fig. 2A, B). In order to more clearly visualize the groupings among the other samples, we also visualized the PCA with the Indian and U13 sample removed. In this PCA, three distinct groups are apparent–Mongolia and Russia; Kyrgyzstan and captive; and Tajikistan, Afghanistan and Pakistan (Fig. 2C).

**Figure 2.**
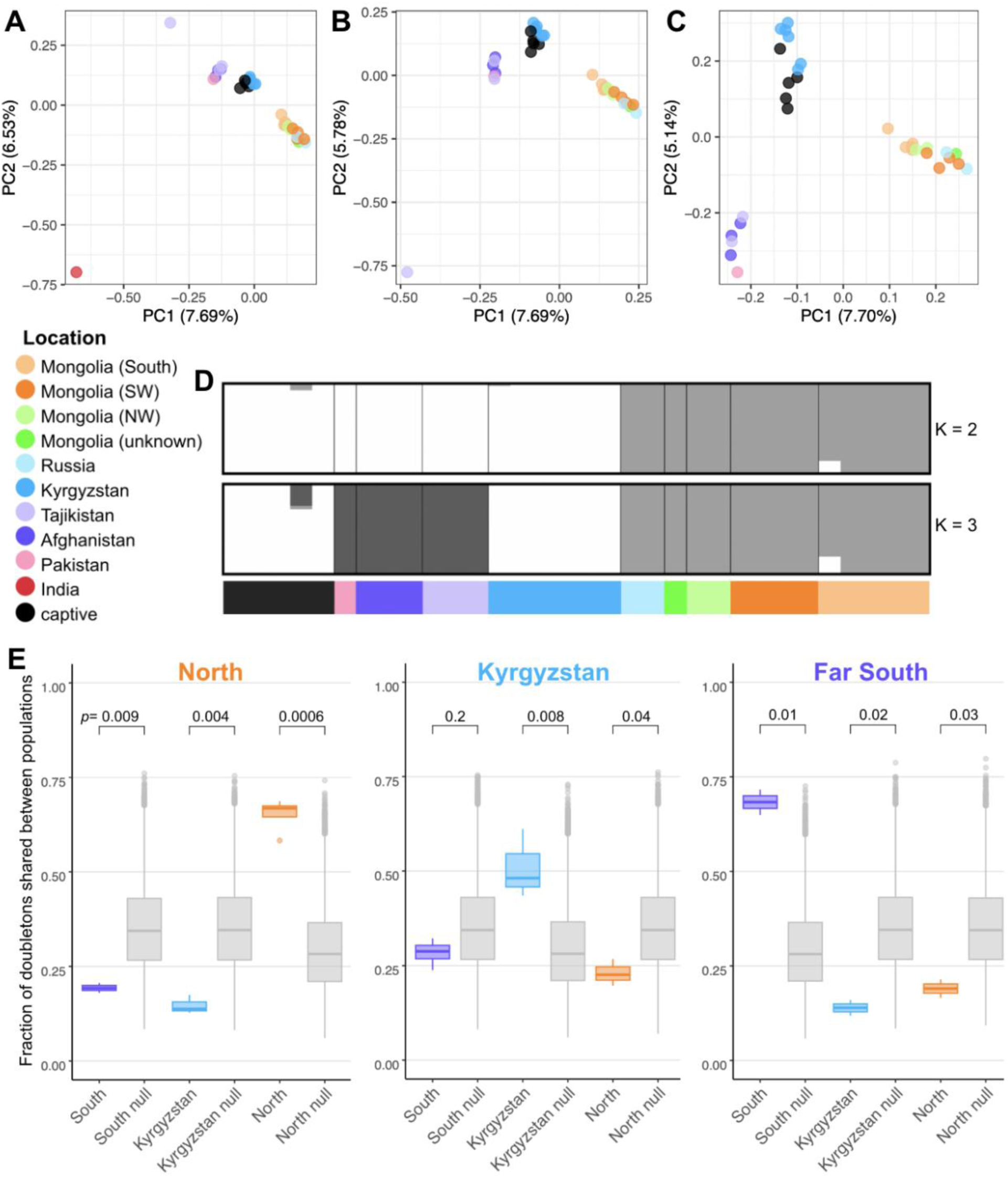
Principal components analysis (PCA), Admixture, and rare allele sharing. A) PCA of genetic variation of 37 unrelated snow leopards using 1,448,657 SNPs. B) PCA after removing one outlier sample from India. C) PCA after removing two outlier samples, India and sample U13 from Tajikistan. PCA axis labels include the percent variation explained by PC1 and PC2. D) Admixture results for ten independent runs for K=2 and K=3 distinct ancestry groups. The ancestry assignments shown for K=2 were supported by all ten iterations and the ancestry assignments shown for K=3 were supported by nine of the ten iterations (the alternate ancestry assignments supported by one run is shown in Supplementary Fig. 1B). E) Doubleton sharing between populations compared to null distributions under panmixia. We identified all doubletons where each allele occurred in a different individual. For each group, identified above each graph, we then calculated the fraction of doubletons occurring in an individual of that group that were shared with individuals of each other group. Observed values are shown in color and null distributions are shown in gray. We made null distributions by randomly shuffling population assignment among the samples and recalculating doubleton sharing 10,000 times. P-values for comparisons between observed data and the null distribution using the Wilcoxon Rank Sum Test are shown. The lower and upper edges of the boxes correspond to the first and third quartiles and the whiskers extend to the lowest/highest value that is no further than 1.5*IQR (inter-quartile range) from the box. Points falling further than 1.5*IQR from the box are plotted individually. In all analyses, we have removed one member of each related pair and in the case of doubleton sharing, we have downsampled populations to n=6 for each group.

We also investigated population structure using Admixture (Alexander et al. 2009) to test models with one to ten ancestral populations (K=1-10) over ten independent runs. Due to issues that can arise with having a small sample size for a genetically distinct group (Nazareno et al. 2017), we ran Admixture analyzes both with and without the India sample (Fig. 2D, Supplementary Fig. 1). Indeed, when including the India sample, Admixture had difficulty assigning it at K=2, with five runs grouping it with the southern samples and five runs showing ancestry split between the north and the south (Supplementary Fig. 1A). Once removing the India sample, Mongolia and Russia separated clearly from all other samples with the same ancestry assignments supported in all ten iterations at K=2. At K=3, also with the India sample removed, nine of the ten iterations supported a clear separation of the samples into three groups–Mongolia and Russia; Tajikistan, Afghanistan, and Pakistan; captives and Kyrgyzstan (Fig. 2D, Supplementary Fig. 1B)–recapitulating what was observed in the PCA. Admixture and PCA results were robust to the removal of captive samples from the analyses (Supplementary Fig. 2).

Maximum likelihood phylogeny construction also showed the Indian sample to be genetically distinct (Supplementary Fig. 3). Thus, PCA, Admixture and phylogenetic results all corroborate that the one Indian sample included in this study is genetically unique from all of the other samples.

Among the other samples (excluding India), Admixture and PCA results suggest three genetically distinct groups. We see a primary genetic divide between the north (Mongolia and Russia) and the south (all other samples) with a secondary divide within the southern group between Kyrgyzstan and populations farther south (Afghanistan, Tajikistan, and Pakistan). These results also show all of our captive samples, whose lineages encompass more than half of all of the founders of the current captive population (33 of 56 founders are represented, Supplementary Fig. 4), group most closely with the Kyrgyzstan samples.

### Assessment of gene flow

Based on Admixture and PCA results, we quantified population structure between the two groups identified in Admixture at K=2 and among the three groups identified at K=3 (Fig. 2D) by assessing shared versus private SNPs, *F_ST_*, and rare allele sharing. We excluded captive samples and the Indian sample from these analyses. We also excluded one sample from each related pair from *F_ST_* and rare allele sharing analyses, but not shared versus private SNP assessments.

At K=2, each group (North and South, down sampled to n=15) had more shared SNPs (598,449 SNPs) than private SNPs (379,861 SNPs private to the north and 364,010 SNPs private to the south) (Supplementary Fig. 5A). At K=3, we compared groups that we will refer to as “North” (consisting of Russia and Mongolia), “Kyrgyzstan”, and “Far South” (consisting of Tajikistan, Afghanistan and Pakistan). We downsampled the North and Far South groups to seven individuals such that all three groups had the same sample size. We found that the Far South and North group had more than twice as many private SNPs (222,569 and 219,175, respectively) as Kyrgyzstan (111,392) (Supplementary Fig. 5B) and that Kyrgyzstan shared a similar number of SNPs with both the North and the Far South (394,020 SNPs shared among all three groups, 65,181 shared only between Kyrgyzstan and North, 64,871 SNPs shared only between Kyrgyzstan and Far South). It is worth noting that the Kyrgyz group represents the smallest geographic area and this limitation in sampling could contribute to our observation of fewer private SNPs in this group.

We calculated Weir and Cockerham’s weighted pairwise *F_ST_*after removing one individual for each first and second degree related pair and downsampling groups to equal size. At K=2, *F_ST_* between the north and south was 0.091. At K=3, the pairwise *F_ST_* between the North and Far South was 0.123, between North and Kyrgyzstan was 0.115, and between Kyrgyzstan and the Far South was 0.093. For context, these values fall well below *F_ST_*observed between different tiger subspecies, which range from 0.164 to 0.318, and are on par with the lower bound *F_ST_* observed between Bengal tiger sub-populations, which range from 0.094 to 0.3 (Armstrong et al. 2021). Observed pairwise *F_ST_*at K=3 was compared to a null distribution representing panmixia created by randomly shuffling population assignments 10,000 times and recalculating *F_ST_*between populations. This analysis showed observed *F_ST_* values to be highly significant (p<0.0001) (Supplementary Fig. 6).

We identified doubletons (SNPs with an alternate allele count of two) where each alternate allele occurs in separate individuals. We then assessed doubleton sharing (Anopheles gambiae 1000 Genomes Consortium et al. 2017) among the three groups–North, Kyrgyzstan, and Far South–after downsampling each group to six individuals. The amount of doubleton sharing between populations can offer a sense of population connectivity as diverged populations have very little rare variant sharing (Gravel et al. 2011). Observed doubleton sharing was compared to a null distribution representing panmixia in the same way as pairwise *F_ST_*–sample population assignments were randomly shuffled and observed doubleton sharing between populations was recalculated 10,000 times. Doubleton sharing corroborated the pre-identified groups with individuals sharing a significantly higher fraction of doubletons (0.40-0.72) with individuals within the same group (p-value North - 0.0006, Kyrgyzstan - 0.008, Far South - 0.01) (Fig. 2E). However, we also found that each individual shared 11-32% of their doubletons with each of the other groups, with the Kyrgyzstan population showing the highest fraction of doubleton sharing with outside groups - even sharing enough with the Far South to not be significantly different from panmixia (p = 0.2). Both Kyrgyzstan and Far South also share enough rare alleles with the North to only be marginally significantly different from panmixia (p=0.04 and p=0.03, respectively). These results support the presence of genetic divergence among these three groups, but also indicate either recent separation or some low level of gene flow among the groups.

### Heterozygosity and historical population size

We used publicly available data to call SNPs in all big cat species using the Genome Analysis Toolkit (GATK) (Van der Auwera and O’Connor 2020) and calculated heterozygosity using VCFtools (Danecek et al. 2011). Here, we defined big cat species as species with an average adult body weight of 40kg or more, which includes all Panthera species (lion (*P. leo*), tiger (*P. tigris*), leopard (*P. pardus*), jaguar (*P. onca*), and snow leopard) as well as cheetah (*Acinonyx jubatus*) and puma (*Puma concolor*) (Johansson et al. 2021). We found snow leopards to have the lowest heterozygosity of any big cat species, with heterozygosity for every snow leopard sample included in this study falling lower than that observed in any other big cat (Fig. 3A). Notably, snow leopard heterozygosity was lower than that of cheetahs, which have long been considered the archetype of low heterozygosity in big cats (Dobrynin et al. 2015; Prost et al. 2022). The relative values of observed heterozygosity among all the other species for which we calculated heterozygosity were consistent with previous work (Paijmans et al. 2021; Pečnerová et al. 2021; Armstrong et al. 2023; Wang et al. 2023).

**Figure 3.**
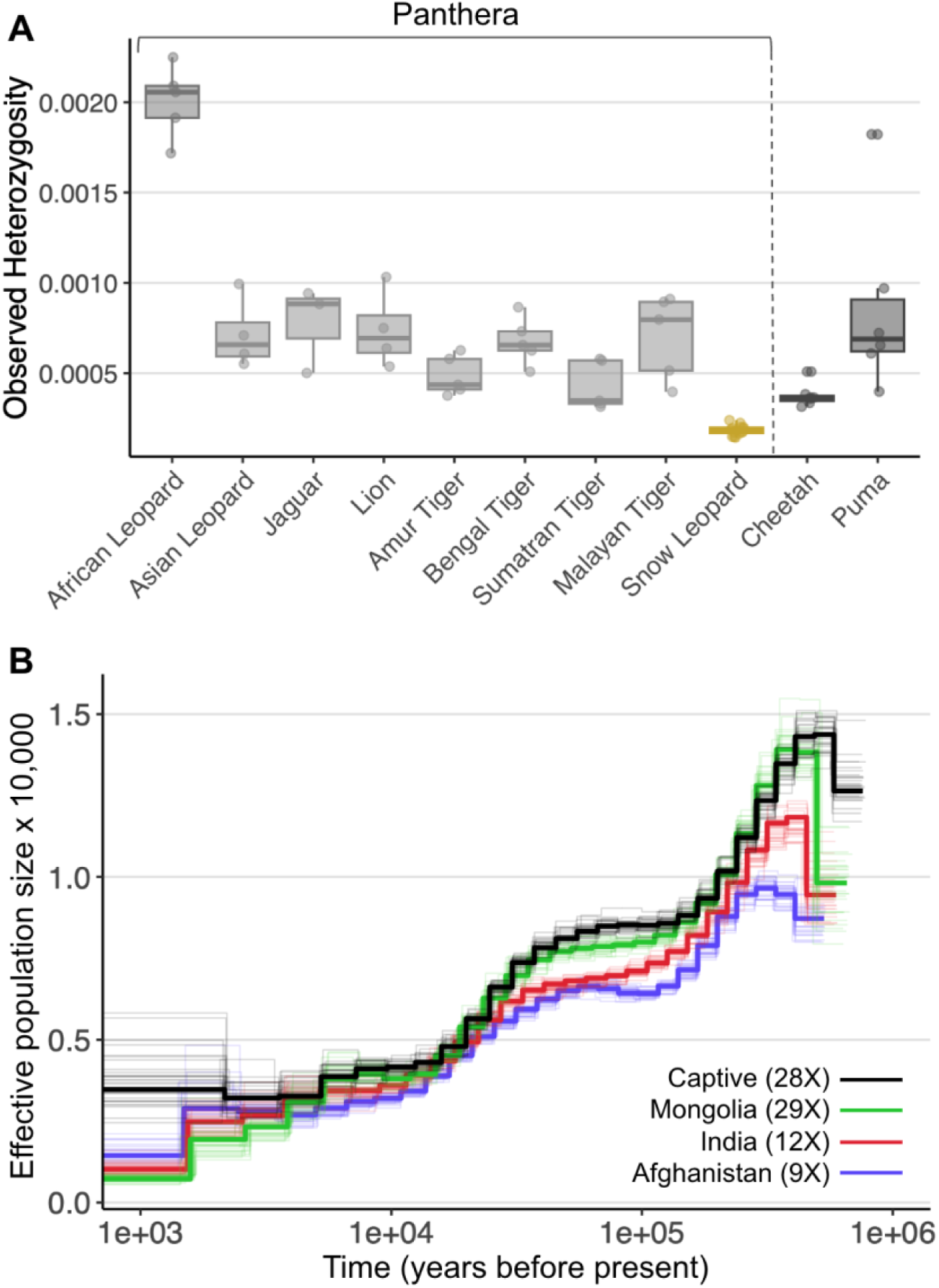
Genome-wide heterozygosity across all big cats and demographic history of snow leopards. A) Comparison of genome-wide heterozygosity across all big cat species. We used publicly available data to call SNPs for every big cat species using GATK and calculated observed heterozygosity using VCFtools. In the case of leopard and tiger, we called SNPs separately for genetically distinct groups. We calculated snow leopard heterozygosity from SNPs called using the same in-house pipeline that was used for SNP calling in all the other big cat species. Only snow leopard samples with a depth of 8X or higher are included (n=15). The lower and upper edges of the boxes correspond to the first and third quartiles and the whiskers extend to the lowest/highest value that is no further than 1.5*IQR (inter-quartile range) from the box. Points falling further than 1.5*IQR from the box are plotted individually. B) Reconstruction of effective population sizes using PSMC using a mutation rate of 0.7×10^-8^ per site per generation and a generation time of five years (Cho et al. 2013). Thirty bootstraps are shown for each sample in thinner, fainter lines of the same color. The average depth of coverage of each sample used is indicated.

We used pairwise sequentially Markovian coalescent (PSMC) (Li and Durbin 2011) to reconstruct historical effective population size using the highest coverage individual from each genetically distinct group (North-29X, Kyrgyzstan/captive-28X, South-9X, India-12X). Reconstructions showed consistent results across all populations sampled and across all bootstrap replicates (Fig. 3B). All reconstructions show a consistently small effective population size over the last ∼300,000 years with snow leopard effective population size never exceeding 14,000 individuals. Reconstructions suggested a snow leopard effective population size of 8,000-6,000 individuals from ∼200,000-20,000 years ago and a decrease to an effective population size of ∼2,500-4,000 individuals ∼20,000 years ago, coincident with the end of the Last Glacial Maximum. Effective population size is generally smaller than census size (Frankham 2007) and PSMC historical effective population size estimates can be impacted by numerous species-specific parameters (e.g., population structure, inbreeding, mutation rate, generation time) (Mazet et al. 2016; Mather et al. 2020) which have not been thoroughly characterized in snow leopards, and thus exact effective population size estimates should be interpreted cautiously.

### Inbreeding and runs of homozygosity (ROH)

Using the same dataset that we used to calculate genome-wide heterozygosity across big cat species, we also calculated the inbreeding coefficient (F) across big cats. These results showed the average inbreeding coefficient of snow leopards to be lower than the puma, cheetah, lion, Asian leopard and numerous tiger subspecies, indicating that the lower genetic diversity observed in snow leopards is not explained by higher inbreeding (Fig. 4A).

**Figure 4.**
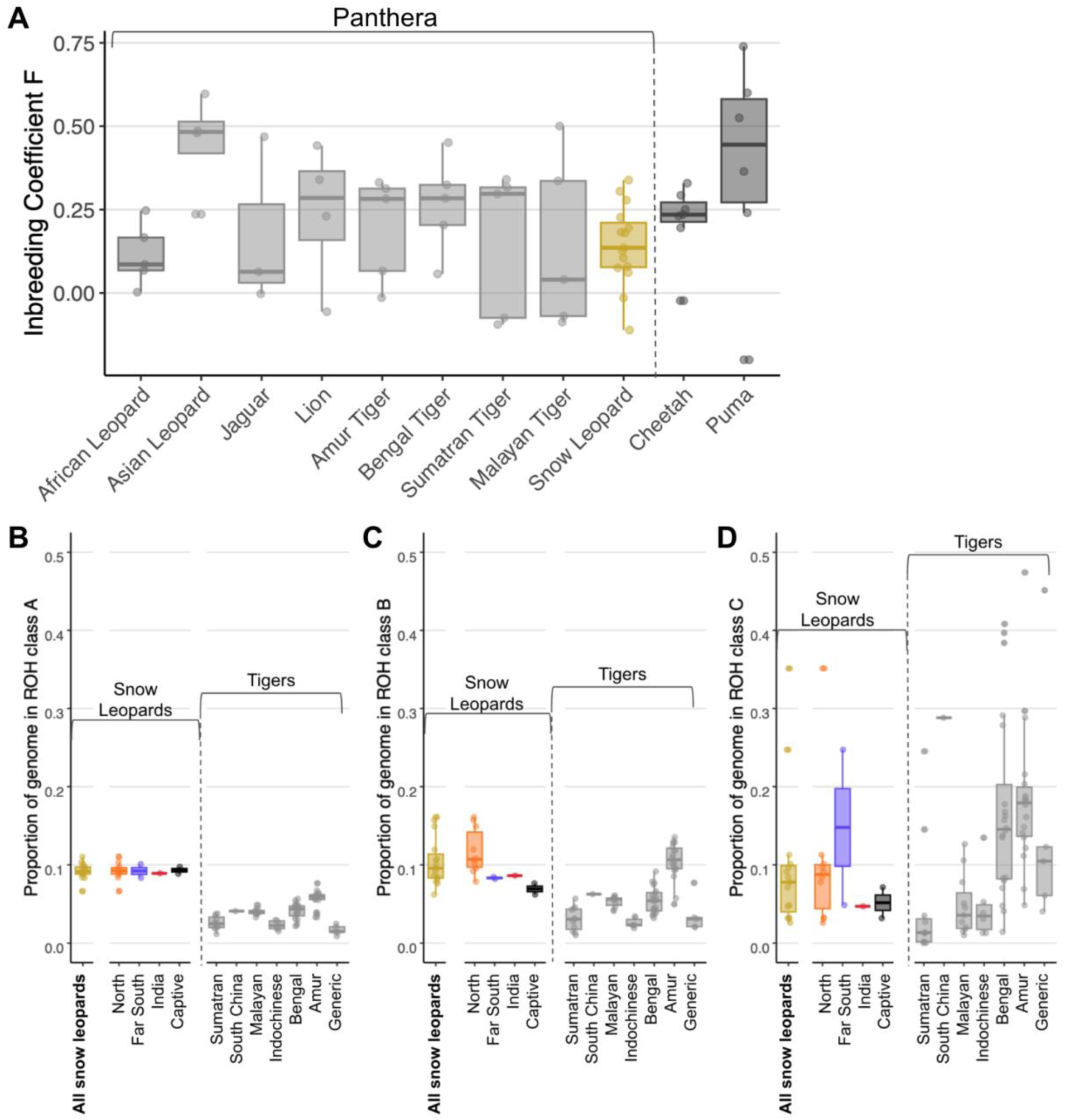
Inbreeding estimates across all big cats and ROH estimates for snow leopards and tigers. A) Comparison of inbreeding coefficient F measured by method of moments across all big cat species. We used publicly available data to call SNPs for every big cat species using GATK and calculated the inbreeding coefficient F using VCFtools. Only snow leopard samples with 8X coverage or greater are included (n=15). B) The proportion of the genome in Class A ROH (small) for snow leopards (colored) and tigers (gray). C) The proportion of the genome in Class B ROH (intermediate) for snow leopards (colored) and tigers (gray). D) The proportion of the genome in Class C ROH (long) for snow leopards (colored) and tigers (gray). All ROH estimates were calculated using GARLIC. We pulled tiger ROH data from Armstrong et al. (Armstrong et al. 2023). Snow leopard ROH is shown separated by geographic group (North, Far South, India, Cpative) as well as all samples together (in gold). Tiger ROH is shown separated by tiger subspecies. All ROH estimates, for snow leopard and tiger, only include samples with greater than 8X coverage. In all plots, each point represents one individual. The lower and upper edges of the boxes correspond to the first and third quartiles and the whiskers extend to the lowest/highest value that is no further than 1.5*IRQ (inter-quartile range) from the box. Points falling further than 1.5*IRQ from the box are plotted individually.

We also used the software GARLIC to assess ROH in snow leopards. GARLIC identifies ROH in three different size classes, each likely the results of different processes–Class A (short) likely reflects homozygosity in ancient ancestral haplotypes, Class B (intermediate) is highly correlated with Class A and indicates recent bottlenecks or background levels of relatedness due to small population size (Pemberton et al. 2012), and Class C (long) reflects recent inbreeding. GARLIC identified 130bp to be the ideal window size for our data set. Garlic uses a sliding window to make an initial logarithm of the odds calculation. Class A was defined as segments under 453,793bp, Class B was defined as segments between 453,793bp - 1,488,700bp, and Class C was defined as segments greater than 1,488,700bp. The proportion of the genome in each class of ROH for each individual was calculated by dividing the total sequence length in each size class by the total mappable length of the genome (1,818,166,894). These proportions were compared to those observed in tigers using GARLIC output provided in Armstrong et al. (Armstrong et al. 2023).

We found the general trends in the proportion of each ROH size class to be different between snow leopards and tigers, which are known to have undergone recent inbreeding due to small population sizes (Khan et al. 2021). On average, snow leopards had a similar proportion of the genome in Class A and B ROH (∼10%) and a little less in Class C ROH (∼8%) with some high outliers. Conversely, in numerous tiger subspecies (Bengal, Amur and Generic), the average proportion of the genome in Class C ROH was much higher than the proportion of the genome in Class A and B.

When comparing ROH trends among snow leopards, we found the proportion of the genome in Class A ROH to be extremely consistent across samples. The proportion of the genome in Class B ROH was also quite consistent across groups, with only a few individuals from the North group showing higher than average values. We observed more variability in the proportion of the genome in Class C ROH across snow leopard samples, with two outlier samples, one from the North and one from the Far South, showing high Class C ROH values.

## Discussion

### Population structure and dispersal barriers

We used WGS from 37 snow leopards to investigate population structure of the species. Our results corroborate many results of previous studies. Population structure analyses show that India is genetically distinct from all other samples (Fig. 2A, Supplementary Fig. 3). The uniqueness of this sample could suggest that this individual is our one representative of the third phylogenetic group (*P. u. uncioides*, Fig. 1B) suggested by Janečka et al. (Janečka et al. 2017), but more samples from this area will be necessary to resolve how genetically distinct Indian snow leopards may be in future analyses.

Among the other samples (excluding India), we identify three genetically distinct groups. Admixture and PCA results identify the most pronounced divide among our samples to occur between the northern and southern part of the range around the Dzungarian Basin (Fig. 2B-D), consistent with previous microsatellite analyses (Janečka et al. 2017; Korablev et al. 2021; Hacker et al. 2023) and models (Riordan et al. 2016; Li et al. 2020). Admixture and PCA results also identify a secondary divide occurring south of Kyrgyzstan (Fig. 2C,D), around the Taklamakan Desert, consistent with previous microsatellite analyses (Korablev et al. 2021). Both of these genetic divides are also supported by highly significant *F_ST_*values between groups (Supplementary Fig. 6).

These results indicate that the Dzungarian Basin and Taklamakan Desert present barriers to dispersal for snow leopards. However, while *F_ST_* values are significant, they are much lower than those observed between tiger subspecies. This, in combination with low levels of differentiation as measured through shared versus private alleles (Supplementary Fig. 5), and shared rare alleles (Fig. 2E) suggest that these genetic divides may be very recent or there may be some level of weak connectivity among these groups. This is consistent with previous fecal microsatellite work which has also found evidence of weak connectivity between Mongolia and China (Hacker et al. 2023), as well as between Kyrgyzstan and Mongolia/Russia and between Kyrgyzstan and Tajikistan (Korablev et al. 2021). Additional samples will be necessary to confidently estimate gene flow in future analyses.

As recommended in Hacker et al. (Hacker et al. 2023), we agree that maintaining this trace level of connectivity should not be prioritized. Rather, connectivity within genetically distinct groups should be targeted especially as human development continues and border fencing, which can potentially act as an absolute barrier to snow leopards, proliferates in this area (Linnell et al. 2016; Li et al. 2020; Johansson et al. 2024).

### Heterozygosity, demographic history and ROH

Our results show that snow leopards have the lowest genetic diversity of any big cat species (Fig. 3A), lower than previously appreciated (Kim et al. 2016; Pečnerová et al. 2021). Demographic history and ROH assessments indicate the exceptionally low heterozygosity in snow leopards is likely due to a persistently small population size over the last 500,000 years (Fig. 3B) rather than from recent inbreeding events. With a persistently small population size and low genetic diversity throughout their evolutionary history, snow leopards serve as yet another example that high genetic diversity is not a requirement for the long-term persistence of a species (Xue et al. 2015; Westbury et al. 2018; Westbury et al. 2019; Morin et al. 2021). Our demographic history assessment (Fig. 3B) did not pick up the more recent population bottleneck ∼8,000 years ago suggested by Janecka et al. (Janečka et al. 2017) from microsatellite data likely because PSMC analyzes loose power at more recent time scales.

Consistent with demographic history assessments, ROH analyses show snow leopards have a greater proportion of their genome in smaller length ROH (Class A and B) (Fig. 4B,C), which likely reflects homozygosity in ancient haplotypes and small historic population size (Pemberton et al. 2012). Conversely, many tiger subspecies, which are known to have high levels of recent inbreeding due to small isolated populations (Khan et al. 2021), show the opposite trend, with the highest proportion of their genome in longer ROH (Class C) (Armstrong et al. 2023) (Fig. 4D). However, we do observe examples of wild snow leopards in the North and the Far South with large portions of their genome in Class C ROH suggesting that there may be sporadic inbreeding events occurring in wild populations.

### Origin of captive samples

Studbook-based pedigrees for the five captive individuals sequenced in this study show that the origin of their wild ancestors is mostly unknown. The few ancestors of known origin are documented as originating from the USSR, Kyrgyzstan, Kazakhstan, and the western Himalayas (Supplementary Fig. 4). Admixture and PCA results both show a strong signal of genetic similarity between the five captive individuals and the current Kyrgyzstan population (Fig. 2A-D). The current Kyrgyzstan population is likely genetically similar to their close neighbors in Kazakhstan and northeastern China, for which we have no genomic data. Thus, these results suggest that the ancestors of the captive individuals largely came from what is currently Kyrgyzstan, or the surrounding area, and fewer ancestors have come from other portions of the range.

### Conservation implications

Snow leopards live in arid, cold, low-productivity, high-elevation habitats where few species can persist – an environment that has evidently only ever been able to support a limited number of snow leopards (Fig. 3B). As a result, they have likely always had an effective population size much lower than other big cats (as suggested in Pečnerová et al.(Pečnerová et al. 2021) and Cho et al. (Cho et al. 2013)), and harbor less genetic diversity than even the cheetah (Fig. 3A).

Thanks to their extreme environment, snow leopards have not yet been exposed to the same level of acute Anthropogenic pressures as have big cats living in habitats more easily accessible to humans. Yet, even having been spared the most intense human impacts, they already have a small population size and extremely low genomic diversity compared to other big cats. While these characteristics have not hindered their long-term persistence thus far, this means that snow leopards do not have the ability to rely on a large population size or standing genetic variation to help them survive any forthcoming Anthropogenic challenges, as other big cats have done (Dinerstein et al. 2007; Bauer et al. 2015). Unfortunately, the most intense Anthropogenic pressures may lie ahead for snow leopards. Anthropogenic climate change threatens to shrink snow leopard range through habitat change (Li et al. 2016) and intensify interspecific competition (e.g. (Pal et al. 2022, Lovari et al. 2024)); shifting grazing practices risk facilitating spillover of novel pathogens from domestic animals into snow leopards and their prey (Ostrowski and Gilbert 2024); and accelerating mining, energy, and infrastructure development threaten to fragment and degrade previously remote snow leopard habitats (Heiner et al. 2024; Zahler and Victurine 2024). Protection of snow leopards and their habitat, to the greatest degree possible, will be pivotal as this low-density carnivore appears genetically and demographically ill-equipped to bounce back from Anthropogenic perturbations (Markert et al. 2010). More broadly, our findings underscore how the IUCN Red List assessment criteria could benefit from the inclusion of genetic factors (Garner et al. 2020) as our results suggest that snow leopards may be more imperiled than their current IUCN Red List status would indicate (Ale and Mishra 2018; Suryawanshi et al. 2019).

## Methods

### Sample collection and sequencing

Through an international collaborative effort, we collected a total of 37 snow leopard blood or tissue samples, composed of 32 wild caught samples from five countries (Afghanistan, Kyrgyzstan, Tajikistan, Mongolia, and Russia), one currently captive but wild-born individual from Pakistan (included in the wild group throughout the manuscript), and four captive samples from mixed/unknown ancestry. Sample details, including DNA extraction and library preparation methods used, can be found in Supplementary Table 1. We sequenced all samples on an Illumina sequencing platform using paired end 150bp reads with an aim of sequencing each sample to ∼5-8X coverage.

All samples from captive individuals were collected as part of routine animal care in the respective AZA zoos and all wild samples were collected as part of ongoing monitoring of snow leopards by local conservation groups with all necessary collection permits. Samples were processed in labs in the United States, France, and Russia and were transported to respective locations with appropriate transport permits.

Additionally, we collected all currently published WGS data for snow leopards, which included data for two captive individuals (NCBI accessions SRR16227515 (Armstrong et al. 2022) and SRR12437590), one wild individual from Mongolia (NCBI accession SRR836372 (Cho et al. 2013)), and one wild individual from India (NCBI BioProject PRJNA1051290). In total, we gathered WGS data for 41 snow leopards.

Note that there is WGS data for an additional sample identified as a snow leopard on Genbank (biosample SAMN17432540, SRA accession SRR13500277, used in the following publications: (Zhou, Wang, et al. 2021; Zhou, Li, et al. 2021)) that we did not use in this study because we concluded that this data was from an Asian leopard (*P. pardus*), not a snow leopard (details of the analyses done to identify this sample as an Asian leopard are provided in the Supplementary Methods and Supplementary Fig. 7).

We were unable to include any samples from a large portion of the snow leopard range in the southeast, but are hopeful that data from this area will be available to include in future analyses.

### Mapping and variant calling

We used the snow leopard reference genome (NCBI accession PRJNA602938 (Armstrong et al. 2022)) and added the lion Y chromosome (from GCA_018350215.1) in order to have the Y-chromosome represented in this female reference genome. After removing duplicate reads, reads were mapped and variants called by Gencove using BWA-MEM (Li 2013) and the GATK (McKenna et al. 2010), respectively. Gencove is a service-providing company that has developed cloud based pipelines to offer quick and streamlined SNP calling. We calculated the depth and breadth of coverage for each sample from bam files using SAMtools (Li et al. 2009) and are provided in Supplementary Table 1.

### Variant filtering

We first filtered variants based on mappability across the genome. We indexed the reference genome and generated mappability scores using GenMap v1.3.0 (Pockrandt et al. 2019) with flags ‘-K 30’ and ‘-E 2’. GenMap mappability scores represent the uniqueness of k-mers (k-mer size given by flag -K) for each position in the genome while allowing for a certain number of mismatches (given by the flag -E) where a mappability score of one at a position indicates that the k-mer at that position occurs only once with up to E mismatches. We sorted the reference genome and mappability bed file using BEDTools (Quinlan and Hall 2010) and removed SNPs falling in a region of the genome with a mappability score less than one using BEDTools *intersect* with the flags ‘-v’, ‘-header’, and ‘-sorted’ (leaving 63,199,070 sites). We then used VCFtools (Danecek et al. 2011) to remove indels using the option ‘--remove-indels’ (leaving 51,199,725 SNPs). We removed individuals with less than 60% of the genome represented (samples U02, U12, and U20) using the VCFtools option ‘--remove-ind’. Most SNPs were removed at this step because sample U02 had over 1.3 million unique singletons (Supplementary Table 1) and over 44 million unique doubletons, likely due to some unknown contamination. We then used VCFtools to remove sites that no longer had any variation using flag ‘--non-ref-ac-any 1’ (leaving 2,379,069 SNPs), SNPs that did not fall on putative autosomes (leaving 2,213,174 SNPs), and non-biallelic SNPs using the flags ‘--min-alleles 2 --max-alleles 2’ (leaving 2,146,294 SNPs). We also split scaffold 22 at base pair 67,650,000 into chromosomes E1 and F1, respectively, to remedy the missassembly described in Armstrong et al.(Armstrong et al. 2022). Next, with bam files as input, we used GATK v4.1.4.1 to add the following annotations to our filtered vcf file using *VariantAnnotator*: QD (QualByDepth - variant confidence normalized by unfiltered depth of variant samples), FS (FisherStrand - strand bias estimated using Fisher’s exact test), MQ (RMSMappingQuality - root mean square of the mapping quality of reads across all samples), MQRankSum (MappingQualityRankSumTest - rank sum test for mapping qualities of reference versus alternate reads), and ReadPosRankSum (ReadPosRankSumTest - rank sum test for relative positioning of reference versus alternate alleles within reads). After adding these annotations, we used GATK *VariantFiltration* to filter SNPs using the following flag: ‘--filter-expression "QD < 2.0 || FS > 60.0 || MQ < 40.0 || MQRankSum < -12.5 || ReadPosRankSum < -8.0"’. We then used VCFtools to remove SNPs that did not pass the GATK *VariantFiltration* step using option ‘--remove-filtered-all’ (leaving 2,065,453 SNPs). Lastly, we removed SNPs missing data in more than 10% of the individuals using flag ‘--max-missing 0.9’ (leaving a final SNP set of 1,591,978). We calculated the number of singletons and private doubletons using the VCFtools flag ‘--singletons’ and removed one of the captive samples (10x_SDzoo) due to an excess of singletons (Supplementary Table 1) which we suspected to be due to the 10X library prep method unique to this sample.

### Relatedness assessment

We estimated relatedness among samples using SNPrelate (Zheng et al. 2012) in R (R Core Team 2020). We divided the samples into three genetically distinct groups based on preliminary PCA and Admixture analyses–Mongolia and Russia, Kyrgyzstan and captive, and all samples South of Kyrgyzstan. We conducted preliminary PCA and Admixture analyses using the same methods described below but in this case included samples that were later removed due to relatedness. We ran each group through SNPrelate separately. We used the function *snpgdsBED2GDS* to convert bed files to gds files. We used *snpgdsLDpruning* to prune the input data using the following flags - ‘method="corr", slide.max.n=50, ld.threshold=0.3, maf = 0.05, autosome.only = F’. We then calculated kinship coefficients using the function *snpgdsIBDMLE* which calculates IBD coefficients for individual pairs using Maximum Likelihood Estimation. A kinship coefficient of 0.5 indicates a duplicate sample, 0.25 indicates a first-order related pair (parent-offspring or full sibling), and 0.125 indicates a second-order related pair (half-siblings or grandparent-grandchild). All sample pairs with non-zero kinship coefficients are listed in Supplementary Table 2. We identified one sample pair with a kinship coefficient greater than that expected from first-order relatives–U01 and U09 (kinship coefficient of 0.335), and we identified three sample pairs with kinship coefficients consistent with second-order related pairs–SL_KGZ_F1 and SL_KGZ_F4 (kinship coefficient of 0.148), AF_SL_07 and AF_SL_06 (kinship coefficient of 0.135), and U14 and U08 (kinship coefficient of 0.124). We removed the sample with lower sequencing coverage from each of these pairs (U01, SL_KGZ_F4, AF_SL_06, and U08) from indicated analyses.

### Pedigrees for captive-bred samples

Using snow leopard studbooks (Blomqvist 2008; Tupa 2011), we compiled information on all of the ancestors of the five captive-born individuals included in this study. The wild origin of their ancestors is mostly unknown; however, each sample has a few ancestors for which there is a wild origin location listed (Supplementary Fig. 4), these origin locations include wild ancestors listed as coming from the USSR (between 1964-1974), Kazakhstan (in 1972), Kyrgyzstan (between 1974-1980), Przewalsk (between 1974-1975, which we believe to refer to Karakol, Kyrgyzstan which was previously named Przewalsk), and Aksai (a contested region between China and India in the Western Himalayas, in 1979).

Using only information from snow leopard studbooks (Blomqvist 2008; Tupa 2011), we visualized pedigrees for each individual with the kinship2 package (Sinnwell et al. 2014) in R (Supplementary Fig. 4).

### Principal component analysis

We conducted principal component analyses (PCAs) on the dataset after filtering for first and second degree relatives (N=33, 1,448,657 SNPs) using PLINK2 (Purcell et al. 2007). First, we generated bed files from vcf files using the flags ‘--make-bed’ and ‘--allow-extra-chr’ and allele frequencies were computed using the flags ‘--freq’ and ‘--allow-extra-chr’. Then, we ran the PCA by inputting the resultant bed file with flag ‘--bfile’ and frequency file with flag ‘--read-freq’ and using the flags ‘--allow-extra-chr’ and ‘--pca 38’. We visualized PCA output using ggplot2 in R.

### Admixture analysis

We conducted Admixture analyses on the dataset after filtering for first and second degree relatives. Since these analyses can be sensitive to having a small sample size for a genetically distinct group (Nazareno et al. 2017), we ran Admixture analyzes both with and without the India sample (Puechmaille 2016; Toyama et al. 2020). We used VCFtools to convert vcf files into PLINK format files. We then used PLINK1.9 to recode alleles numerically using the flags ‘--allow-extra-chr --recode12’. We then ran Admixture (Alexander et al. 2009) for K=1-10 ten times using the flag ‘-s time’ so that a different random seed was used for each run based on the time. We visualized all runs using Clumpak (Kopelman et al. 2015).

We also conducted PCA and Admixture analyses on the dataset after removing captive samples in order to assess the impact of captive samples on population structure results (Supplementary Fig. 2) as we know these individuals have at least somewhat mixed ancestry and have been separated from the wild population for many decades.

### Phylogenetic tree construction

We constructed a phylogenetic tree as an additional way to visualize genetic groupings among the samples using the same data set used for PCA and Admixture analyses. We included two tiger samples (SRR13647578 and SRR13500748) to serve as an outgroup. We first mapped the tiger WGS data to the snow leopard reference genome using BWA-MEM. We then used the function *vcf2bed* from bedops (Neph et al. 2012) with the flags ‘-snvs’ and ‘-d’ to convert our filtered snow leopard vcf file to a bed file. We then used BCFtools *mpileup* with the flags ‘-A -a AD,DP -R’ followed by BCFtools *call* with the flag ‘-m’ to generate a vcf file of SNP calls for just the snow leopard SNP sites from the tiger data. We then merged the tiger calls to our snow leopard vcf file using *vcf-merge* from vcftools. We used vcf2phylip (Creators Edgardo M. Ortiz1 Show affiliations 1. Technical University of Munich) to convert the merged vcf to phylip format and constructed a maximum likelihood phylogeny using IQtree (Nguyen et al. 2015) with the flags ‘-st DNA -m GTR+ASC -T 10 -B 1000’. We visualized the tree using FigTree v1.4.4 (Anon).

### Quantifying population structure

We further characterized population divides identified in Admixture and PCA analyses by calculating the number of shared versus private SNPs among groups, pairwise *F_ST_*, and the rate of rare variant sharing among groups. We excluded captive samples from these analyses as well as the sample from India as the PCA indicates that this sample is genetically distinct from all groups.

We did not filter for relatedness when calculating shared versus private SNPs among the groups; however, we did downsample as necessary to have an equal number of samples in each group for all comparisons (n=15 for the K=2 north vs. south comparison and n=7 for the K=3 North-Kyrgyzstan-FarSouth comparison). We used BCFtools (Li 2011) to create and index vcf files with only the individuals from each group using the *view* and *index* function, respectively. We then used BCFtools *isec* to calculate how many SNPs were shared and private among the groups.

Before calculating *F_ST_*, we removed one individual for each first or second degree related pair. We also downsampled to have an equal number of samples in each group (n=13 for the K=2 north vs. south comparison and n=6 for the K=3 North-Kyrgyzstan-FarSouth comparison). We then used vcftools to calculate the Weir and Cockerham weighted *F_ST_* estimate (Weir and Cockerham 1984) between each group using the flag ‘--weir-fst-pop’. In order to assess the significance of *F_ST_* estimates at K=3, we conducted 10,000 permutations by shuffling the population assignments and recalculating *F_ST_* to generate a null distribution.

To assess doubleton sharing among groups, we excluded one individual for each related pair and downsampled all groups so that each had an equal number of individuals (N=6). We then used VCFtools to retain only SNPs with a minor allele count of two (doubletons) using the flags ‘--mac 2 --max-mac 2’. We removed any SNPs where the doubletons were present in the same individual and identified allele sharing among samples using the flag ‘--genome full’ in PLINK. We created a null distribution under panmixia by calculating doubleton sharing between populations after randomly shuffling population assignments among the samples 10,000 times. We compared fractions of observed doubleton sharing to the null distribution using the Wilcoxon Rank Sum Test in R.

### PSMC

We used PSMC (Li and Durbin 2011) to estimate snow leopard effective population size back in time using the highest coverage sample for each population cluster. We generated diploid consensus sequences for each individual as recommended by PSMC. Starting with bam files filtered to include only putative autosomes, we used SAMtools *mpileup* and BCFtools *call* to generate a vcf. We then used *vcfutils.pl vcf2fq* to generate diploid fastq files using the ‘-D’ and ‘- d’ flags to reflect the average depth of coverage of each sample as recommended in PSMC documentation. We then ran PSMC with the default settings. We also performed 30 rounds of bootstrapping using random sampling with replacement for each sample. For plotting, we used a mutation rate of 0.7×10^-8^ per site per generation as suggested for snow leopards by Cho et al. (Cho et al. 2013) and a generation time of five years.

### Heterozygosity in other big cats

We calculated heterozygosity in all big cat species using publicly available data in order to see how snow leopard heterozygosity levels compare to other big cat species. We included all other species in the genus *Panthera* (leopard, lion, tiger, jaguar) as well as cheetah and puma. Accession numbers of publicly available WGS data and reference genomes used in this analysis are listed in Supplementary Table 3 and 4, respectively. We mapped all fastq data to the corresponding reference genome using BWA-MEM (Li 2013). We sorted and indexed resulting bam files using SAMtools (Li et al. 2009). We added read groups and marked duplicates using picard (Anon). We calculated depth of coverage and breadth of coverage for each sample using SAMtools. We then used GATK (Van der Auwera and O’Connor 2020) *HaplotypeCaller* to generate a gvcf file for each sample and GATK *CombineGVCFs* to combine the gvcf files for every species (or subgroup in the case of tigers and leopards). Lastly, we used GATK *GenotypeGVCFs* to create a final vcf file for every group.

In order to limit the need for excessive computing power, we downsampled some samples with over 30X coverage using SAMtools *view* as indicated in Supplementary Table 3. The leopard samples required a larger amount of computing power than the other species to call haplotypes, so we split these samples into one bam file per contig using BAMtools *split* and ran each contig through GATK *HaplotypeCaller* and *CombineGVCFs* in parallel. We also found it necessary to split the jaguar and snow leopard data into intervals to combine gvcf files using GATK *GenomicsDBImport* followed by *CombineGVCFs*. In both cases, we concatenated resultant vcf files, one per contig or interval, using BCFtools *concat*.

We filtered each reference genome for mappability using genmapv1.3.0 to first index and then calculate mappability using flags ‘-K 30’ and ‘-E 2’. We then filtered each vcf to only include nucleotides with a mappability score of one. To do this, we sorted both vcf files and mappability bed files using BEDTools and retained only SNPs falling in regions with a mappability score of one in each vcf using BEDTools *intersect*. We then filtered each vcf using GATK *VariantFiltration* with the following flag: ‘--filter-expression "QD < 2.0 || FS > 60.0 || MQ < 40.0 || MQRankSum < -12.5 || ReadPosRankSum < -8.0"’. Then, we filtered vcf files to remove indels and non-bialellic SNPs using VCFtools flags ‘--remove-indels --min-alleles 2 --max-alleles 2’.

We calculated the mappable length of each genome using mappability bed files to calculate the number of nucleotides with a mappability score of one. Using the filtered vcf for each species or group, we calculated observed homozygosity for each sample using VCFtools (Danecek et al. 2011) with the flag ‘--het’. In R, we calculated the number of heterozygous sites by subtracting the number of observed homozygous sites column ((O)HOM) from the total number of sites column (NSITES). We then calculated heterozygosity by dividing the number of heterozygous sites by the length of the genome consisting of putative autosomes with a mappability score of one. We used ggplot2 in R to create box plots of heterozygosity results.

### Heterozygosity in snow leopards

Although SNPs had already been called by Gencove (as described above), we re-called SNPs from the snow leopard dataset using the same in-house pipeline used to call SNPs in all of the other big cat species to ensure comparability of heterozygosity calculations among species. We calculated observed heterozygosity in snow leopards using both SNP datasets (Gencove’s and ours) in the same way described above. We calculated heterozygosity for all individuals, regardless of relatedness (N=37); however, because heterozygous SNPs are most accurately called with higher coverage data (Maruki and Lynch 2017), we only used samples with 8X coverage or more.

We compared snow leopard heterozygosity calculated from SNPs called using our in-house pipeline to heterozygosity calculated from SNPs called by Gencove using Pearson correlation coefficient calculated using the ggpubr (Kassambara 2020) package in R (Supplementary Fig. 8). We found heterozygosity calculated from the two different SNP calling pipelines to be extremely correlated (R = 0.97, p=5.1E-9); however, all heterozygosity estimates calculated from our in-house SNP calls were slightly higher (average of ∼0.000056) than that estimated from Gencove SNP calls.

### Runs of homozygosity

We converted vcf files to transposed PLINK files using plink version 1.90b5.3 using the flags ‘--allow-extra-chr –const-fid 0 --recode transpose’. We then inferred ROH in each individual using the software GARLIC (Szpiech et al. 2017) with an error rate of 0.001 and the centromere location of each chromosome set to 0, 0; since centromere location is unknown. In addition, we used ‘--auto-win-size, --auto-overlap-frac, –winsize 100’ to allow GARLIC to determine the ideal window size to use. We excluded runs of homozygosity that were less than 50,000bp.

ROH were automatically divided into different size classes by GARLIC - A (short), B (intermediate), and C (long). We calculated the proportion of the genome in each size class by dividing the total sequence length in each size class by the total sequence length of putative autosomes with a mappability score greater than one. The proportion of the genome in each size class was visualized in box plots using ggplot2 in R.

We pulled ROH values for tigers, calculated using GARLIC, from Armstrong et al. (Armstrong et al. 2023) and calculated the proportion of the genome in each size class of ROH for each individual in the same way as described above, but using the total mappable length across all autosomes in the tiger reference genome.

As with heterozygosity assessments in snow leopards, in order to limit any biases caused by allelic drop out in lower coverage samples, we only used samples with 8X coverage or higher in ROH assessments in both snow leopards and tigers.

### Inbreeding across big cats

We calculated the inbreeding coefficient F in all big cat species using the same SNP datasets used to calculate heterozygosity. As with heterozygosity assessments, we only calculated inbreeding for snow leopards samples with 8X coverage or higher. We calculated the inbreeding coefficient F using the method of moments ((observed homozygous count - expected count)/(total observations - expected count)) using vcftools with the flag ‘-het’.

### Samples included in each analysis

We provide a detailed list of which samples were included in each analysis in Supplementary Table 5.

## Data Availability

Data associated with this study has been deposited into bioproject PRJNA1048427 and will be released upon publication.

## Code Availability

The code used for analyses in this project is available on the project’s github: https://github.com/ksolari/SL_WGS

## Acknowledgements

This work was supported by funding from the Snow Leopard Trust. We would like to thank those who helped us in acquiring the captive samples used in this study, including Karen Ingerman and Susie Bartlett from the Bronx Zoo. We would like to thank Haiqing Xu for his help in conducting simulations to produce the null distribution for the shared doubleton analysis.

