## Supplementary Materials for "Extreme in Every Way: Exceedingly Low Genetic Diversity in Snow Leopards Due to Persistently Small Population Size"

### Supplementary Methods:

**Species ID of Biosample SAMN17432540:**

Publicly available whole-genome sequencing data was downloaded from NCBI for at least two individuals from each species in the Panthera clade - lion (ERR4139877, ERR4139880), tiger (SRR13647584, SRR13647599), jaguar (SRR4444359, SRR11097154), leopard (Asian - ERR5671313, ERR5671317, SRR5382750; African - ERR5671315, ERR5671314) and snow leopard (SRR13500277, SRR836372). We used BWA-MEM[1] to map all of this data, as well as whole-genome sequencing data from five additional snow leopard samples that we generated, to the snow leopard reference genome[2]. Mapped data was sorted and indexed using SAMtools[3]. We then used ANGSD[4] followed by ngstools[5] to generate a PCA. A total of 40,205,810 variable sites were identified in ANGSD. All of the Panthera species group clearly in the PCA except for biosample SAMN17432540 (SRR13500277) which clearly groups with Asian leopards and not snow leopards (Supplementary Figure 7).

#### Supplementary Figures:

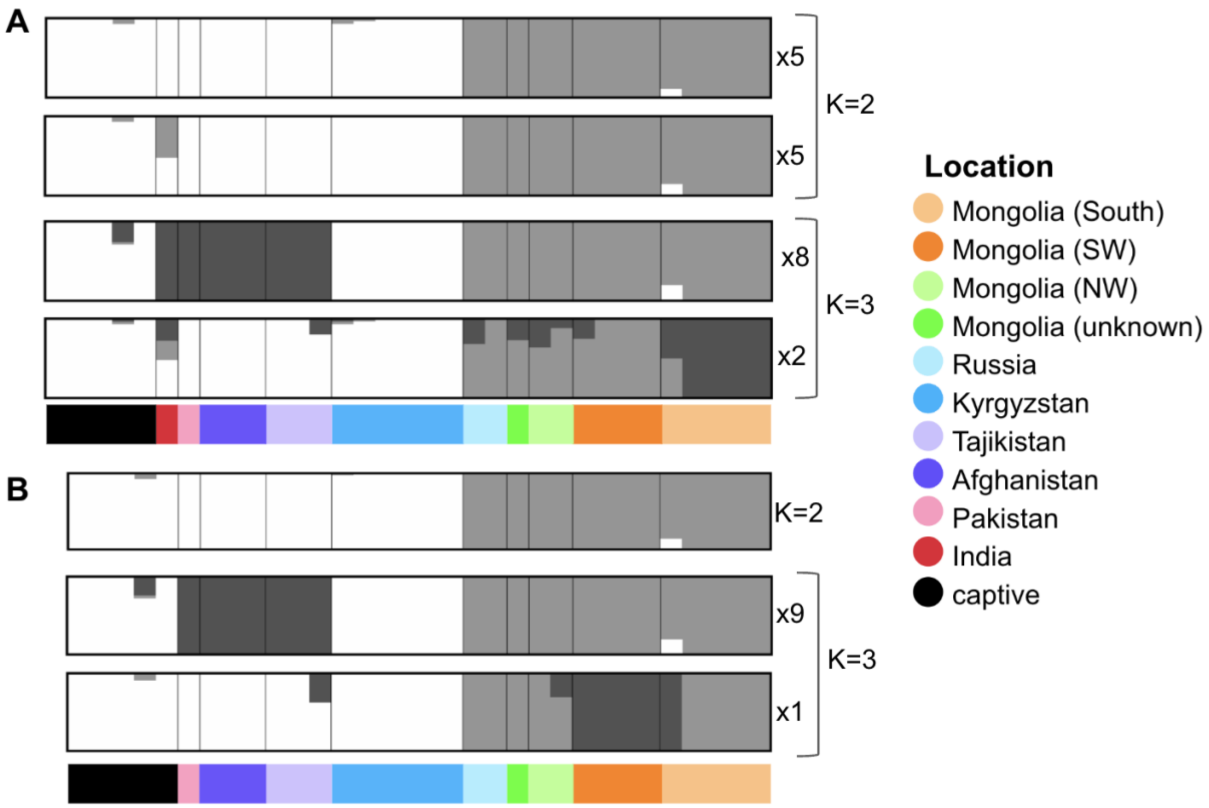


**Supplementary Figure 1. Admixture results with and without the India sample.** A) Admixture results for ten independent runs for K=2 and K=3 distinct ancestry groups with all non-related individuals included (n=37). The number to the right of each bar plot indicates the number of iterations supporting each. B) Admixture results for ten independent runs for K=2 and K=3 distinct ancestry groups with the India sample excluded. The ancestry assignments shown for K=2 were supported by all ten iterations and the ancestry assignments shown for K=3 were supported by nine (top) and one (bottom) of the ten iterations. The top two admixture plots shown in B are also included in Figure 2 of the main manuscript.

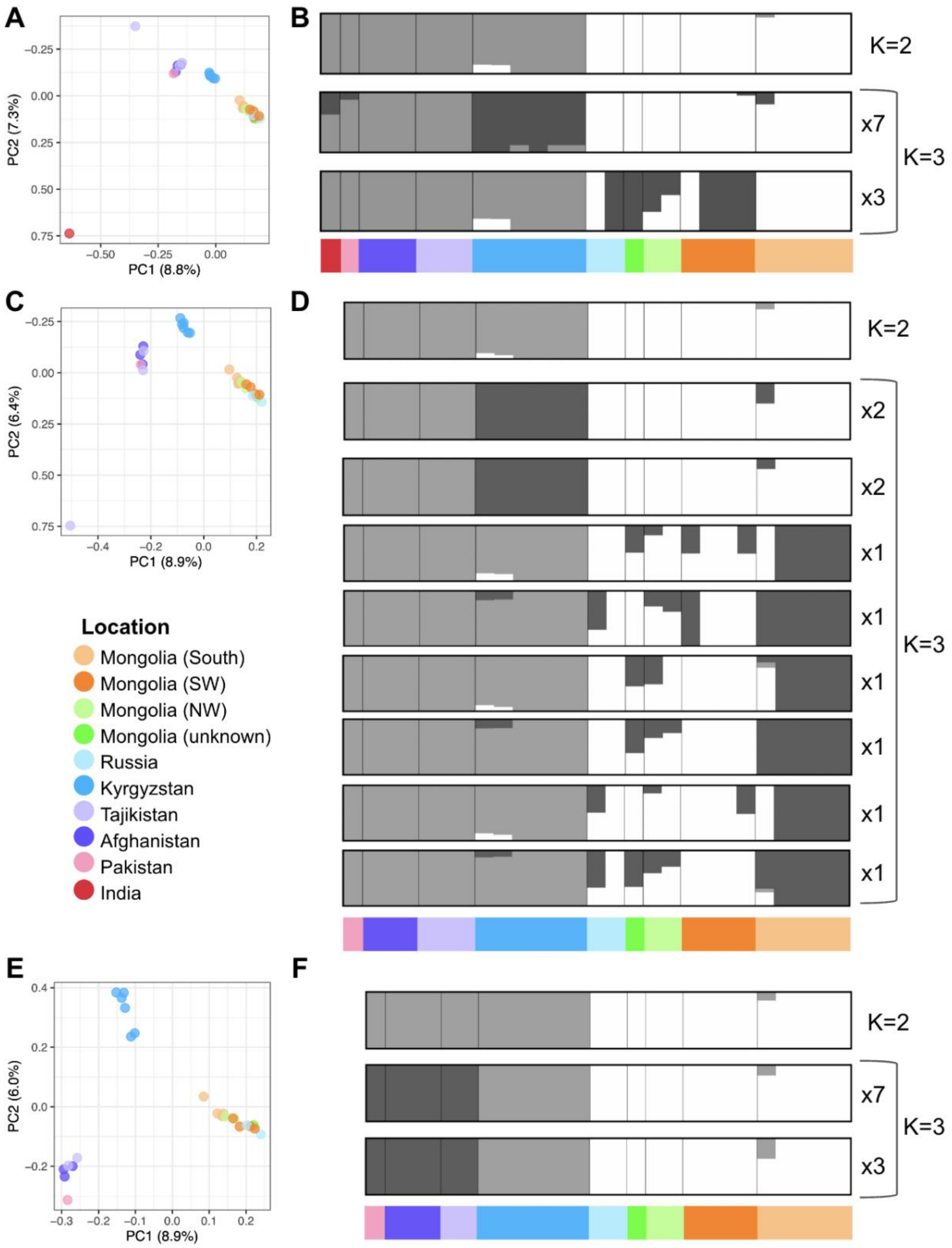


**Supplementary Figure 2. Admixture and PCA results with captive samples removed.** A)

PCA of genetic variation for all non-related wild-origin samples. B) Admixture results for all non-related wild-origin samples. C) PCA of genetic variation for all non-related wild-origin samples excluding India. D) Admixture results for all non-related wild-origin samples excluding India. E) PCA of genetic variation for all non-related wild-origin samples excluding India and one Tajikistan sample, U13. F) Admixture results for all non-related wild-origin samples excluding India and one Tajikistan sample, U13. PCA axis labels include the percent variation explained by PC1 and PC2. All Admixture plots show ten independent runs for K=3 and K=2. In all cases, the ancestry assignments shown for K=2 are supported by all ten iterations. The number of iterations supporting each K=3 Admixture plot is indicated to the right of each.

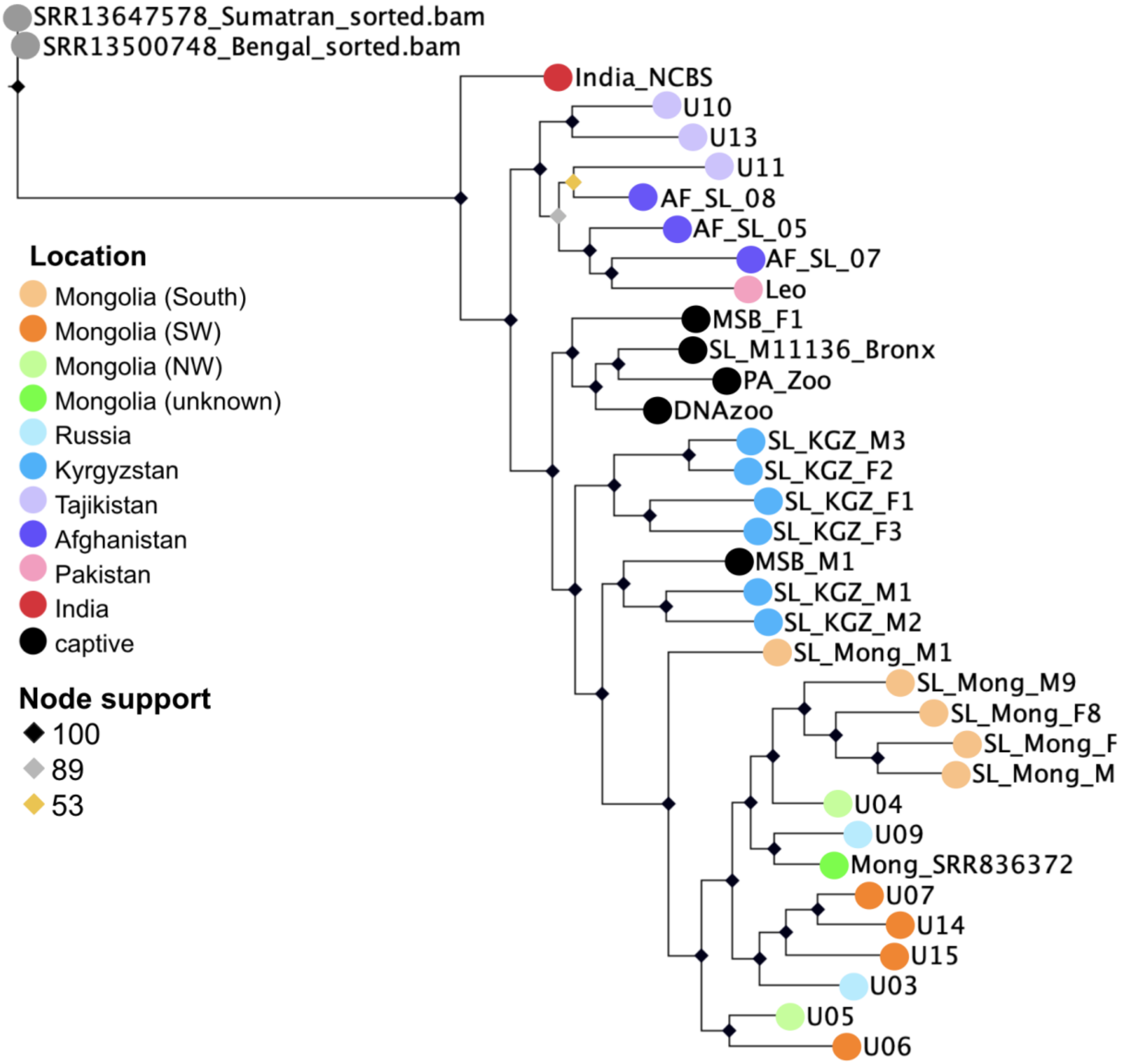


**Supplementary Figure 3. Snow leopard phylogeny.** Maximum likelihood phylogeny using genome-wide SNP data from unrelated snow leopard samples with two tiger samples serving as an outgroup. 1000 bootstraps were run and node support is indicated by different colored diamonds.

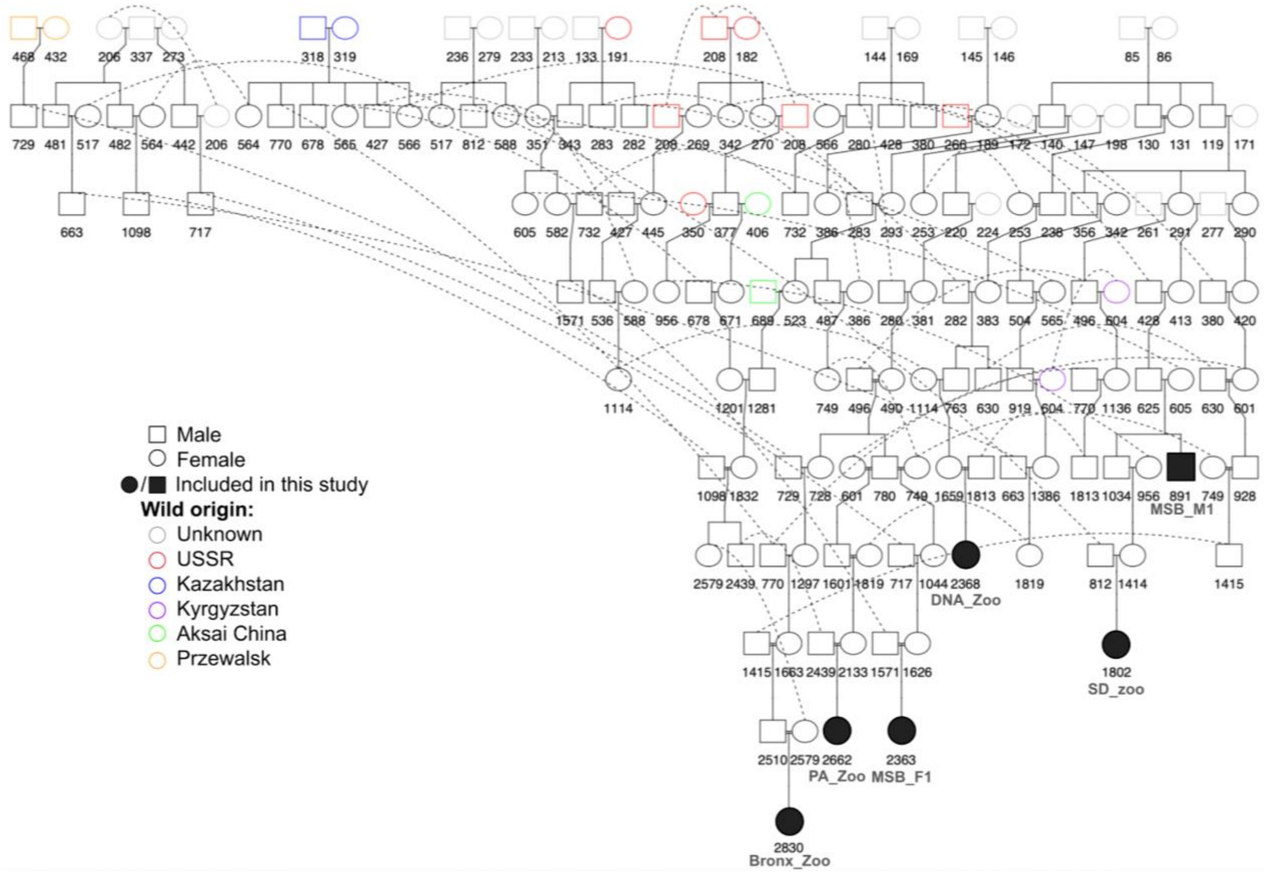


**Supplementary Figure 4. Pedigree of captive snow leopards included in this study.** This pedigree was constructed using data from studbooks and visualized using the Kinship2 package in R. The origin of wild founders are indicated with different colors as identified in the legend. All individuals shown with a black outline of a circle (female) or square (male) are captive-born. Studbook numbers are shown under each individual. Individuals included in this study are shown with solid black shapes and sample names are provided under studbook numbers.

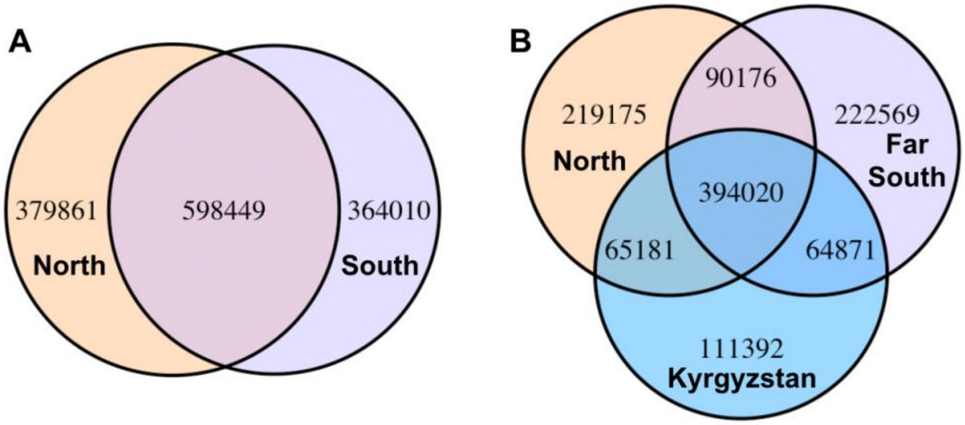


**Supplementary Figure 5. Shared and private SNPs among sample locations.** A) Comparing North (N=15) versus South (N=15) as identified by Admixture at K=2 (1,342,320 SNPs total). B) Comparing North, Kyrgyzstan, and Far South (all subsampled to n=7) as identified by Admixture at K=3 (1,167,384 SNPs total). Captive individuals and the Indian sample are not included in these analyses.

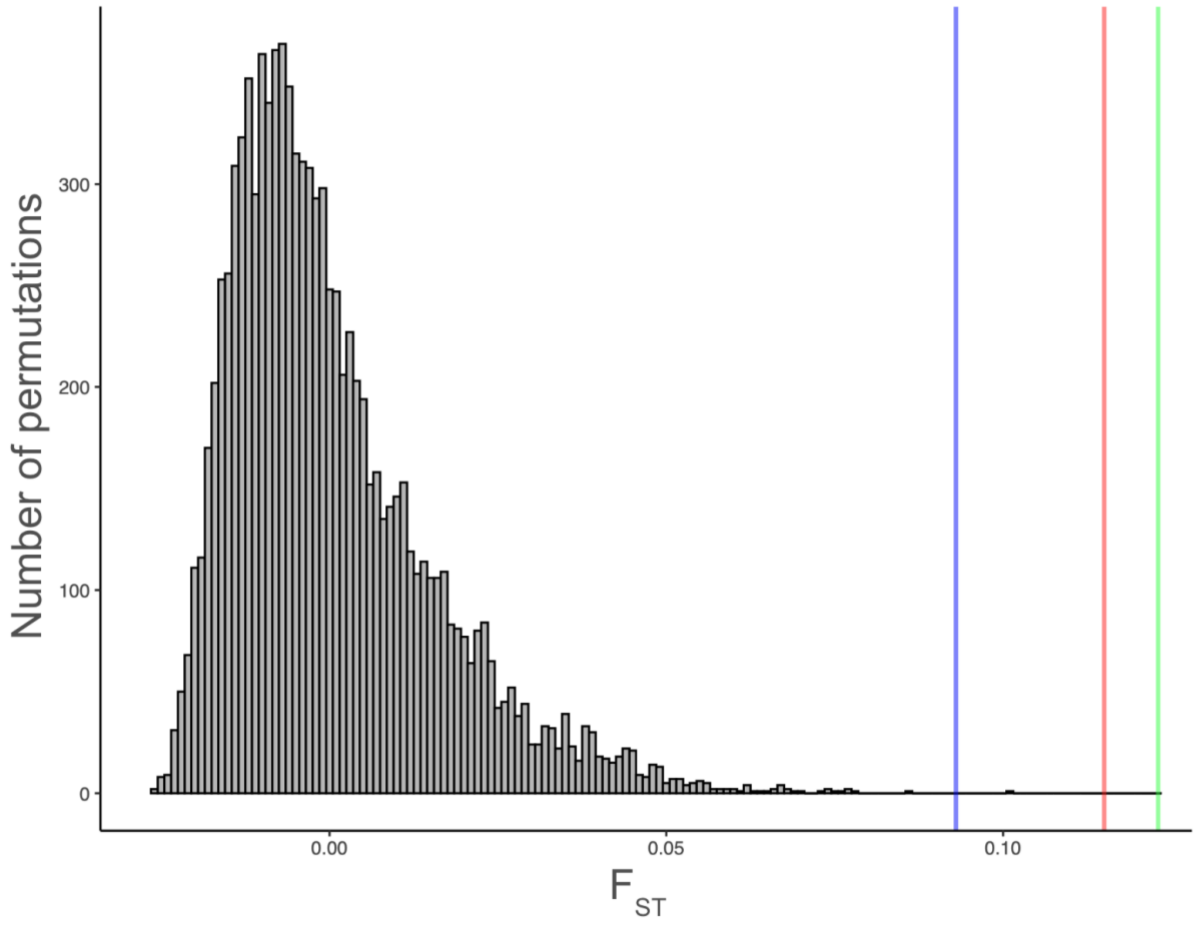


**Supplementary Figure 6. Comparison of observed $F_{ST}$ values to the null distribution.** The null distribution of weighted pairwise $F_{ST}$ generated by randomly permuting individual location assignments 10,000 times is shown in grey. Observed $F_{ST}$ between the North and Far South (0.123) is shown with a vertical green line. Observed $F_{ST}$ between North and Kyrgyzstan (0.115) is shown with a vertical red line. Observed $F_{ST}$ between Kyrgyzstan (0.093) and the Far South is shown with a vertical blue line.

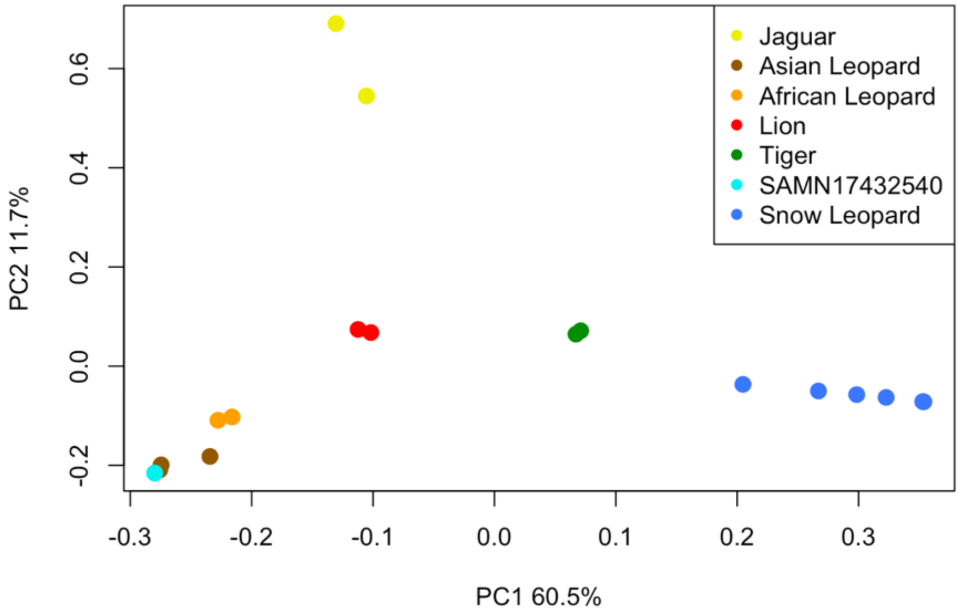


**Supplementary Figure 7. Principal component analysis showing the species ID of Biosample SAMN17432540.** The PCA is based on whole-genome sequencing data. PC1 and PC2 are visualized and the percent variance explained by each principal component is indicated.

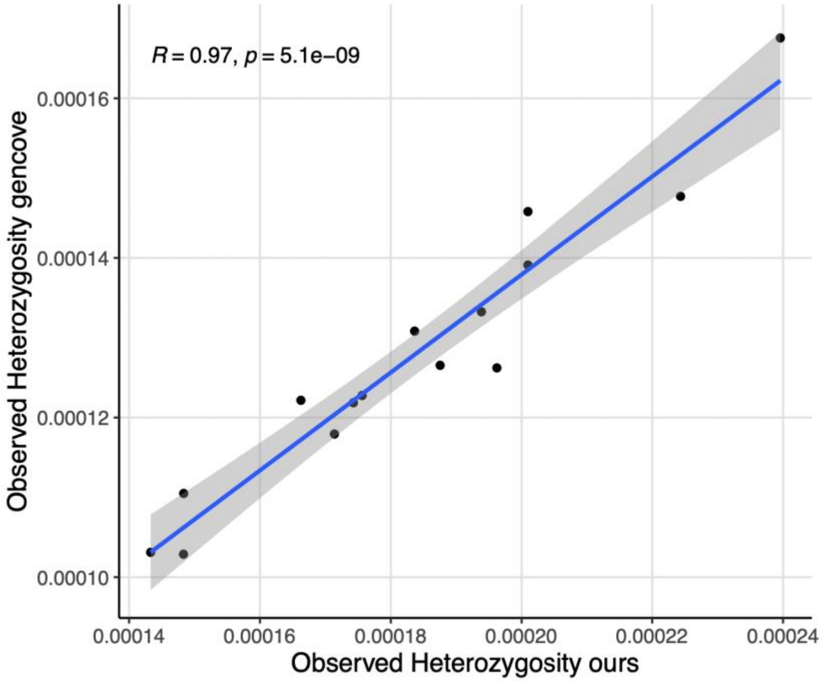


**Supplementary Figure 8. Comparison of observed heterozygosity calculated from different implementations of GATK to call SNPs.** Observed heterozygosity calculated using SNPs called with our in-house pipeline are shown on the x-axis and heterozygosity calculated using SNPs called by Gencove are shown on the y-axis. Pearson correlation coefficient calculated using the ggpubr package in R is indicated.

#### Supplementary Tables:

##### Supplementary Table 1. Sample Information.

| Sample ID | Country | Source | Depth | Breadth | # of singletons | Stud book # | Lat | Long | Collection year | Sex | NCBI Accession | DNA extraction method | Library preparation method |
|---|---|---|---|---|---|---|---|---|---|---|---|---|---|
| SD_zoo* | captive | Armstrong et al., 2022 | 33.65 | 99.1 | 81952 | 1802 | | | | F | SAMN13911153 | Qiagen MagAttract kit | 10× Genomics Chromium library preparation |
| DNA_Zoo | captive | DNA zoo | 28.37 | 99.03 | 12253 | 2368 | | | | F | SAMN15801464 | | |
| MSB_F1 | captive | current study | 5.37 | 95.72 | 4090 | 2363 | | | | F | SAMN38638818 | Qiagen DNeasy Blood and Tissue kit | Nextera DNA Library Prep |
| MSB_M1 | captive | current study | 5.37 | 96.08 | 4491 | 891 | | | | M | SAMN38638819 | Qiagen DNeasy Blood and Tissue kit | Nextera DNA Library Prep |
| PA_Zoo | captive | current study | 4.66 | 94.84 | 4036 | 2662 | | | | F | SAMN38638820 | Qiagen DNeasy Blood and Tissue kit | Nextera DNA Library Prep |
| Bronx_Zoo | captive | current study | 8.12 | 98.11 | 11983 | 2830 | | | | F | SAMN38638821 | Qiagen DNeasy Blood and Tissue kit | Nextera DNA Library Prep |
| Mong_F8 | Mongolia (South) | current study | 5.7 | 95.68 | 3059 | | 43.199 | 100.436 | 2012 | F | SAMN38638822 | Qiagen DNeasy Blood and Tissue kit | Nextera DNA Library Prep |
| Mong_F9 | Mongolia (South) | current study | 5.37 | 96.16 | 2900 | | 43.272 | 100.707 | 2012 | F | SAMN38638823 | Qiagen DNeasy Blood and Tissue kit | Nextera DNA Library Prep |
| Mong_M1 | Mongolia (South) | current study | 6.22 | 96.37 | 7574 | | 43.134 | 100.550 | 2008 | M | SAMN38638824 | Qiagen DNeasy Blood and Tissue kit | Nextera DNA Library Prep |
| Mong_M10 | Mongolia (South) | current study | 5.3 | 95.85 | 2834 | | 43.183 | 100.437 | 2012 | M | SAMN38638825 | Qiagen DNeasy Blood and Tissue kit | Nextera DNA Library Prep |
| Mong_M9 | Mongolia (South) | current study | 6.88 | 97.07 | 3634 | | 43.151 | 100.694 | 2011 | M | SAMN38638826 | Qiagen DNeasy Blood and Tissue kit | Nextera DNA Library Prep |
| U06 | Mongolia (SW) | current study | 10.72 | 95.61 | 17607 | | 47.796 | 90.872 | 2015 | M | SAMN38638827 | Isogen Diatom DNA kit | NEBNext Ultra II DNA Library Prep |
| U07 | Mongolia (SW) | current study | 11.65 | 98.6 | 5082 | | 47.569 | 92.587 | 2015 | F | SAMN38638828 | Qiagen QIAmp Fast DNA Tissue Kit | NEBNext Ultra II DNA Library Prep |

| | | | | | | | | | | | | | |
|---|---|---|---|---|---|---|---|---|---|---|---|---|---|
| U08 | Mongolia (SW) | current study | 9.19 | 98.43 | 2627 | | 47.561 | 92.591 | 2015 | M | SAMN38638829 | Qiagen QIAmp Fast DNA Tissue Kit | NEBNext Ultra II DNA Library Prep |
| U14 | Mongolia (SW) | current study | 10.86 | 98.84 | 1648 | | 46.673 | 93.441 | 2017 | M | SAMN38638830 | Qiagen QIAmp Fast DNA Tissue Kit | NEBNext Ultra II DNA Library Prep |
| U15 | Mongolia (SW) | current study | 4.71 | 90.35 | 3044 | | 46.445 | 93.463 | 2016 | F | SAMN38638831 | Isogen Diatom DNA kit | NEBNext Ultra II DNA Library Prep |
| U20* | Mongolia (SW) | current study | 1.37 | 46.59 | 10663 | | 47.906 | 90.872 | 2019 | | SAMN38638832 | CTAB[7] | NEBNext Ultra II DNA Library Prep |
| U04 | Mongolia (NW) | current study | 9.57 | 98.56 | 4474 | | 50.294 | 91.068 | 2014 | F | SAMN38638833 | CTAB[7] | NEBNext Ultra II DNA Library Prep |
| U05 | Mongolia (NW) | current study | 16.79 | 99.03 | 4099 | | 50.286 | 91.119 | 2015 | M | SAMN38638834 | Qiagen QIAmp Fast DNA Tissue Kit | NEBNext Ultra II DNA Library Prep |
| Mong_SRR836372 | Mongolia (unknown) | Cho et al., 2013 | 28.87 | 98.86 | 14363 | | | | | F | SAMN02086968 | | |
| U01 | Russia | current study | 9.59 | 98.12 | 10550 | | 51.300 | 89.787 | 2009 | | SAMN38638835 | CTAB[7] | NEBNext Ultra II DNA Library Prep |
| U02* | Russia | current study | 5.14 | 53.73 | 1322527 | | 51.847 | 92.068 | 2013 | M | SAMN38638836 | CTAB[7] | NEBNext Ultra II DNA Library Prep |
| U03 | Russia | current study | 9.96 | 97.74 | 21605 | | 50.737 | 86.900 | 2015 | F | SAMN38638837 | Qiagen QIAmp Fast DNA Tissue Kit | NEBNext Ultra II DNA Library Prep |
| U09 | Russia | current study | 9.61 | 98.45 | 10453 | | 50.784 | 89.740 | 2013 | | SAMN38638838 | CTAB[7] | NEBNext Ultra II DNA Library Prep |
| KGZ_F1 | Kyrgyzstan | current study | 5.06 | 93.87 | 3963 | | 41.945 | 78.576 | 2015 | F | SAMN38638839 | Qiagen DNeasy Blood and Tissue kit | Nextera DNA Library Prep |
| KGZ_F2 | Kyrgyzstan | current study | 4.64 | 93.4 | 3443 | | 41.977 | 78.537 | 2016 | F | SAMN38638840 | Qiagen DNeasy Blood and Tissue kit | Nextera DNA Library Prep |
| KGZ_F3 | Kyrgyzstan | current study | 3.73 | 89.93 | 3603 | | 41.910 | 78.591 | 2017 | F | SAMN38638841 | Qiagen DNeasy Blood and Tissue kit | Nextera DNA Library Prep |
| KGZ_F4 | Kyrgyzstan | current study | 4.48 | 93.07 | 2654 | | 41.909 | 78.590 | 2017 | F | SAMN38638842 | Qiagen DNeasy Blood and Tissue kit | Nextera DNA Library Prep |
| KGZ_M1 | Kyrgyzstan | current study | 6.14 | 95.87 | 5193 | | 41.873 | 78.579 | 2016 | M | SAMN38638843 | Qiagen DNeasy Blood and Tissue kit | Nextera DNA Library Prep |

| | | | | | | | | | | | | | |
|---|---|---|---|---|---|---|---|---|---|---|---|---|---|
| KGZ_M2 | Kyrgyzstan | current study | 4.78 | 93.47 | 4115 | | 41.909 | 78.590 | 2016 | M | SAMN38638844 | Qiagen DNeasy Blood and Tissue kit | Nextera DNA Library Prep |
| KGZ_M3 | Kyrgyzstan | current study | 6.27 | 96.17 | 5150 | | 41.876 | 78.578 | 2016 | M | SAMN38638845 | Qiagen DNeasy Blood and Tissue kit | Nextera DNA Library Prep |
| U10 | Tajikistan | current study | 6.56 | 88.07 | 7407 | | 37.875 | 73.691 | 2016 | F | SAMN38638846 | Isogen Diatom DNA kit | NEBNext Ultra II DNA Library Prep |
| U11 | Tajikistan | current study | 7.3 | 73.44 | 4185 | | 37.875 | 73.692 | 2016 | | SAMN38638847 | Isogen Diatom DNA kit | NEBNext Ultra II DNA Library Prep |
| U12* | Tajikistan | current study | 1 | 13.62 | 1319 | | 37.835 | 74.076 | 2018 | | SAMN38638848 | CTAB[7] | NEBNext Ultra II DNA Library Prep |
| U13 | Tajikistan | current study | 6.71 | 95.15 | 22190 | | 38.455 | 74.372 | 2018 | | SAMN38638849 | CTAB[7] | NEBNext Ultra II DNA Library Prep |
| AF_05 | Afghanistan | current study | 7.66 | 98.5 | 10554 | | 38.301 | 71.177 | 2019 | F | SAMN38638850 | Macherey-Nagel Nucleospin Genomic DNA from tissue | Illumina DNA Prep |
| AF_06 | Afghanistan | current study | 3 | 85.4 | 6779 | | 38.200 | 70.933 | 2020 | F | SAMN38638851 | Macherey-Nagel Nucleospin Genomic DNA from tissue | Illumina DNA Prep |
| AF_07 | Afghanistan | current study | 3.4 | 87.73 | 8081 | | 36.948 | 72.973 | 2010 | | SAMN38638852 | Macherey-Nagel Nucleospin Genomic DNA from tissue | Illumina DNA Prep |
| AF_08 | Afghanistan | current study | 9.25 | 98.83 | 10463 | | 36.963 | 72.902 | 2011 | | SAMN38638853 | Macherey-Nagel Nucleospin Genomic DNA from tissue | Illumina DNA Prep |
| Leo | Pakistan (wild born, currently captive) | current study | 9.06 | 98.13 | 8484 | 2630 | 36.249 | 74.045 | 2021 | M | SAMN38638854 | Qiagen DNeasy Blood and Tissue kit | Nextera DNA Library Prep |
| India_NCBS | India | National Centre for Biological Sciences-TIFR | 12.01 | 98.88 | 49158 | | 32.300 | 78.010 | 2012 | | PRJNA1051290 | | |

*Samples that were removed from all analyses due to sequencing quality issues.

**Supplementary Table 2. Kinship coefficients calculated using SNPrelate for all pairs with non-zero coefficients.**

| Group | Individual 1 | Individual 2 | Kinship |
|---|---|---|---|
| North | U09 | U01* | 0.33499001 |
| Kyrgyzstan_captive | SL_KGZ_F1 | SL_KGZ_F4* | 0.14820347 |
| Far South | AF_SL_07 | AF_SL_06* | 0.13538874 |
| North | U14 | U08* | 0.12415888 |
| Kyrgyzstan_captive | SL_KGZ_F2 | SL_KGZ_M3 | 0.06992244 |
| North | U08 | U07 | 0.06411454 |
| North | SL_Mong_M10 | SL_Mong_F9 | 0.0628048 |
| North | U14 | U07 | 0.06229824 |
| North | SL_Mong_M10 | SL_Mong_F8 | 0.04878393 |
| North | Mong_SRR836372 | U04 | 0.03749832 |
| Kyrgyzstan_captive | SL_KGZ_F3 | SL_KGZ_F4 | 0.0343196 |
| Kyrgyzstan_captive | SL_KGZ_M2 | SL_KGZ_M1 | 0.03340161 |
| North | SL_Mong_F9 | SL_Mong_F8 | 0.03131735 |
| North | U15 | U07 | 0.02262996 |
| North | SL_Mong_M10 | SL_Mong_M9 | 0.02188175 |
| North | Mong_SRR836372 | U05 | 0.01539328 |
| North | SL_Mong_F9 | SL_Mong_M9 | 0.01527608 |
| Kyrgyzstan_captive | PA_Zoo | SL_M11136_Bronx | 0.01430921 |
| Kyrgyzstan_captive | SL_KGZ_F3 | SL_KGZ_F1 | 0.01388335 |
| Kyrgyzstan_captive | DNAzoo | SL_M11136_Bronx | 0.0096397 |
| North | U03 | U07 | 0.00852837 |
| North | U05 | U04 | 0.00745974 |
| North | SL_Mong_F8 | SL_Mong_M9 | 0.00723796 |
| Kyrgyzstan_captive | MSB_F1 | MSB_M1 | 0.00595172 |
| North | U06 | SL_Mong_M1 | 0.00470249 |
| North | SL_Mong_M1 | U06 | 0.00470249 |
| North | U15 | U14 | 0.00460162 |
| North | U03 | U04 | 0.00440541 |
| North | U03 | U06 | 0.00404295 |
| Kyrgyzstan_captive | SL_KGZ_F2 | SL_KGZ_F3 | 0.00282453 |
| North | U08 | U05 | 0.0023464 |
| North | U15 | U06 | 0.00190692 |
| Kyrgyzstan_captive | SL_M11136_Bronx | MSB_F1 | 0.00167512 |

*Samples removed from analyses potentially impacted by the presence of related individuals as indicated in the main text.

**Supplementary Table 3. Published whole-genome sequencing data used for heterozygosity comparisons among all big cat species.**

| **Species/group** | **sample** | **depth** | **breadth** | **Observed Heterozygosity** |
|---|---|---|---|---|
| Amur Tiger | SRR13647652 | 5.81 | 96.27 | 0.000376324 |
| Amur Tiger | SRR13647654 | 7.70 | 98.68 | 0.000410507 |
| Amur Tiger | SRS8209282 | 17.46 | 99.45 | 0.000578811 |
| Amur Tiger | SRS8209284 | 13.89 | 99.32 | 0.000626396 |
| Amur Tiger | SRS8209286 | 8.67 | 99.10 | 0.000435949 |
| Bengal Tiger | SRR13500746 | 8.45 | 99.32 | 0.000655783 |
| Bengal Tiger | SRR13500748 | 15.01 | 99.41 | 0.000865245 |
| Bengal Tiger | SRR13500752 | 21.20 | 99.49 | 0.000625797 |
| Bengal Tiger | SRR13500756 | 18.19 | 99.53 | 0.000508425 |
| Bengal Tiger | SRR13500762 | 8.95 | 99.37 | 0.000731831 |
| Malayan Tiger | SRR13647621 | 6.20 | 97.63 | 0.000396791 |
| Malayan Tiger | SRR13647627 | 6.14 | 97.10 | 0.000514727 |
| Malayan Tiger | SRS8209294 | 16.35 | 99.43 | 0.000796851 |
| Malayan Tiger | SRS8209296 | 32.27 | 99.56 | 0.000910584 |
| Malayan Tiger | SRS8209300 | 22.27 | 99.53 | 0.000894811 |
| Sumatran Tiger | SRR13647577 | 6.35 | 97.52 | 0.000347958 |
| Sumatran Tiger | SRR13647578 | 6.32 | 97.50 | 0.000331245 |
| Sumatran Tiger | SRR13647591 | 5.67 | 96.56 | 0.000314467 |
| Sumatran Tiger | SRS8209305 | 31.23 | 99.52 | 0.00057024 |
| Sumatran Tiger | SRS8209313 | 22.27 | 99.54 | 0.000578745 |
| Lion | SRR10009886 | 37.66 | 99.15 | 0.001032482 |
| Lion | SRR13242484 | 21.22 | 98.98 | 0.000749692 |
| Lion | SRR836361 | 29.38 | 98.77 | 0.000638026 |
| Lion | SRR836370 | 24.16 | 98.70 | 0.00053714 |
| Jaguar | SRR11097154 | 22.75* | 99.00 | 0.000883412 |
| Jaguar | SRR14572000 | 21.67* | 98.66 | 0.000500771 |
| Jaguar | SRR4444360 | 22.42* | 98.52 | 0.000941729 |
| African Leopard | ERR5671299 | 19.50 | 99.23 | 0.002090843 |

| | | | | |
|---|---|---|---|---|
| African Leopard | ERR5671307 | 16.42 | 98.98 | 0.002248533 |
| African Leopard | ERR5671308 | 18.04* | 99.35 | 0.00205636 |
| African Leopard | ERR5671314 | 20.22* | 99.85 | 0.001718174 |
| African Leopard | ERR5671315 | 22.64 | 99.69 | 0.001914143 |
| Asian Leopard | ERR5671303 | 12.55 | 81.12 | 0.000606921 |
| Asian Leopard | ERR5671311 | 18.14* | 99.30 | 0.000551796 |
| Asian Leopard | ERR5671313 | 22.08* | 99.48 | 0.000709197 |
| Asian Leopard | ERR5671323 | 7.87 | 96.00 | 0.000994117 |
| Cheetah | SRR2737540 | 5.64 | 97.58 | 0.000314751 |
| Cheetah | SRR2737541 | 7.12 | 98.23 | 0.000358603 |
| Cheetah | SRR2737542 | 7.76 | 98.19 | 0.000365219 |
| Cheetah | SRR2737543 | 6.64 | 98.10 | 0.00036498 |
| Cheetah | SRR2737544 | 6.29 | 97.81 | 0.000335003 |
| Cheetah | SRR2737545 | 7.52 | 98.13 | 0.000383392 |
| Cheetah | SRR9951918 | 24.29 | 99.55 | 0.000508819 |
| Puma | BR406 ( SRR7542886, SRR7542887, SRR7542888) | 32.49 | 94.10 | 0.001822242 |
| Puma | CYP47 (SRR7664677, SRR7664678) | 42.56 | 94.18 | 0.000398165 |
| Puma | EVG21 (SRR7660678, SRR7660679) | 51.92 | 94.23 | 0.000655054 |
| Puma | SC29 (SRR7537344, SRR7537345) | 29.22 | 94.15 | 0.000722349 |
| Puma | SMM22 (SRR7543017, SRR7543018) | 34.81 | 94.18 | 0.000609408 |
| Puma | YNP198 (SRR7610940, SRR7610941) | 39.03 | 94.19 | 0.00097003 |

In all cases, the whole-genome sequencing data was mapped to the reference genome for that species (Supplementary Table 4), SNPs were called using GATK, and heterozygosity was estimated using VCFtools. *Bam files that were down sampled. The depth after down sampling is shown.

**Supplementary Table 4. Genome mappability calculations for all big cat reference genomes used in heterozygosity analyses.**

| Species | Reference genome | Total bp | unmappable bp | mappable bp |
|---|---|---|---|---|
| Lion | GCA_008795835.1 | 2387958570 | 441792892 | 1946165678 |
| Puma | GCF_003327715.1 | 2314358962 | 401315200 | 1913043762 |
| Jaguar | GCA_028533385.1 | 2468400402 | 544548340 | 1923852062 |
| Tiger | GCA_021130815.1 | 2405965077 | 460360448 | 1945604629 |
| Cheetah | GCF_003709585.1 | 2373450562 | 428515259 | 1944935303 |
| Leopard | GCA_024362965.1 | 2441143628 | 494445445 | 1946698183 |
| Snow Leopard | GCF_023721935.1 | 2205089538 | 386922644 | 1818166894 |

**Supplementary Table 5. Summary of which samples were included in each analysis and why certain samples were removed.**

| | | | **Analysis** | PCA | PCA | PCA | PCA | PCA | PCA | Admixture | Admixture | Admixture | Admixture | Admixture | Phylogeny | Shared vs. Private SNPs at K=2 | Shared vs. Private SNPs at K=3 | $F_{ST}$ for K=2 | $F_{ST}$ for K=3 | Rare variant sharing at K=3 | Genome-wide heterozygosity | Inbreeding | ROH |
|---|---|---|---|---|---|---|---|---|---|---|---|---|---|---|---|---|---|---|---|---|---|---|---|
| | | | **Figure** | 2A | 2B | 2C | S2A | S2B | S2C | S1A | 2D, S1B | S2B | S2D | S2F | S3 | S5A | S5B | | | 2E | 3A | 4A | 4B,C,B |
| | | | **Sample Removal Criteria** | Relatives | Relatives, India | Relatives, India, U13 | Relatives, captives | Relatives, India, captives | Relatives, India, U13, captives | Relatives | Relatives, India | Relatives, captives | Relatives, India, captives | Relatives, India, captives, U13 | Relatives | Captives, India, down sampled to n=15 for each group | Captives, India , down sampled to n=7 for each group | Relatives, captives, India, down sampled to n=13 for each group | Relatives, captives, India, down sampled to n=6 for each group | Relatives, captives, India, down sampled to n=6 for each group | Samples with less than 8X coverage | Samples with less than 8X coverage | Samples with less than 8X coverage |
| **Sample ID** | **Location** | **Depth** | | | | | | | | | | | | | | | | | | | | | |
| DNA_Zoo | captive | 28.37 | | X | X | X | | | | X | X | | | | X | | | | | | X | X | X |
| MSB_F1 | captive | 5.37 | | X | X | X | | | | X | X | | | | X | | | | | | | | |
| MSB_M1 | captive | 5.37 | | X | X | X | | | | X | X | | | | X | | | | | | | | |

| | | | | | | | | | | | | | | | | | | | | | | | |
|---|---|---|---|---|---|---|---|---|---|---|---|---|---|---|---|---|---|---|---|---|---|---|---|
| PA_Zoo | captive | 4.66 | | X | X | X | | | | X | X | | | | X | | | | | | | | |
| Bronx_Zoo | captive | 8.12 | | X | X | X | | | | X | X | | | | X | | | | | | X | X | X |
| Mong_F8 | Mongolia (South) | 5.7 | | X | X | X | X | X | X | X | X | X | X | X | X | X | X | X | X | X | | | |
| Mong_F9 | Mongolia (South) | 5.37 | | X | X | X | X | X | X | X | X | X | X | X | X | X | X | X | | | | | |
| Mong_M1 | Mongolia (South) | 6.22 | | X | X | X | X | X | X | X | X | X | X | X | X | X | | X | | | | | |
| Mong_M10 | Mongolia (South) | 5.3 | | X | X | X | X | X | X | X | X | X | X | X | X | X | | | | | | | |
| Mong_M9 | Mongolia (South) | 6.88 | | X | X | X | X | X | X | X | X | X | X | X | X | X | | X | | | | | |
| U06 | Mongolia (SW) | 10.72 | | X | X | X | X | X | X | X | X | X | X | X | X | X | | X | | | X | X | X |
| U07 | Mongolia (SW) | 11.65 | | X | X | X | X | X | X | X | X | X | X | X | X | X | X | X | X | X | X | X | X |
| U08 | Mongolia (SW) | 9.19 | | | | | | | | | | | | | | X | | | | | X | X | X |
| U14 | Mongolia (SW) | 10.86 | | X | X | X | X | X | X | X | X | X | X | X | X | X | | X | | | X | X | X |
| U15 | Mongolia (SW) | 4.71 | | X | X | X | X | X | X | X | X | X | X | X | X | X | | X | | | | | |
| U04 | Mongolia (NW) | 9.57 | | X | X | X | X | X | X | X | X | X | X | X | X | X | X | X | X | X | X | X | X |
| U05 | Mongolia (NW) | 16.79 | | X | X | X | X | X | X | X | X | X | X | X | X | X | X | X | X | X | X | X | X |
| Mong_SRR 836372 | Mongolia (unknown) | 28.87 | | X | X | X | X | X | X | X | X | X | X | X | X | X | | X | | | X | X | X |
| U01 | Russia | 9.59 | | | | | | | | | | | | | | | | | | | X | X | X |
| U03 | Russia | 9.96 | | X | X | X | X | X | X | X | X | X | X | X | X | X | X | X | X | X | X | X | X |
| U09 | Russia | 9.61 | | X | X | X | X | X | X | X | X | X | X | X | X | X | X | X | X | X | X | X | X |
| KGZ_F1 | Kyrgyzstan | 5.06 | | X | X | X | X | X | X | X | X | X | X | X | X | X | X | X | X | X | | | |
| KGZ_F2 | Kyrgyzstan | 4.64 | | X | X | X | X | X | X | X | X | X | X | X | X | X | X | X | X | X | | | |
| KGZ_F3 | Kyrgyzstan | 3.73 | | X | X | X | X | X | X | X | X | X | X | X | X | X | X | X | X | X | | | |

| | | | | | | | | | | | | | | | | | | | | | | | |
|---|---|---|---|---|---|---|---|---|---|---|---|---|---|---|---|---|---|---|---|---|---|---|---|
| KGZ_F4 | Kyrgyzstan | 4.48 | | | | | | | | | | | | | | X | X | | | | | | |
| KGZ_M1 | Kyrgyzstan | 6.14 | | X | X | X | X | X | X | X | X | X | X | X | X | X | X | X | X | X | | | |
| KGZ_M2 | Kyrgyzstan | 4.78 | | X | X | X | X | X | X | X | X | X | X | X | X | X | X | X | X | X | | | |
| KGZ_M3 | Kyrgyzstan | 6.27 | | X | X | X | X | X | X | X | X | X | X | X | X | X | X | X | X | X | | | |
| U10 | Tajikistan | 6.56 | | X | X | X | X | X | X | X | X | X | X | X | X | X | X | X | X | X | | | |
| U11 | Tajikistan | 7.3 | | X | X | X | X | X | X | X | X | X | X | X | X | X | X | X | X | X | | | |
| U13 | Tajikistan | 6.71 | | X | X | | X | X | | X | X | X | X | | X | X | X | X | X | X | | | |
| AF_05 | Afghanistan | 7.66 | | X | X | X | X | X | X | X | X | X | X | X | X | X | X | X | X | X | | | |
| AF_06 | Afghanistan | 3 | | | | | | | | | | | | | | X | | | | | | | |
| AF_07 | Afghanistan | 3.4 | | X | X | X | X | X | X | X | X | X | X | X | X | X | X | X | | | | | |
| AF_08 | Afghanistan | 9.25 | | X | X | X | X | X | X | X | X | X | X | X | X | X | X | X | X | X | X | X | X |
| Leo | Pakistan (wild born, currently captive) | 9.06 | | X | X | X | X | X | X | X | X | X | X | X | X | X | X | X | X | X | X | X | X |
| India_NCBS | India | 12.01 | | X | | | X | | | X | | X | | | X | | | | | | X | X | X |

Note that samples U02, U12, U20 and SD_zoo were removed from all analyses for sample quality issues described in the methods. Xs indicate samples that were included in each analysis. In the case of analyses where we subsampled to have an equal number of samples from each group, samples included are colored by group membership.
